# Quantitative analyses of T cell motion in tissue reveals factors driving T cell search in tissues

**DOI:** 10.1101/2022.11.17.516891

**Authors:** David J Torres, Paulus Mrass, Janie Byrum, Arrick Gonzales, Dominick Martinez, Evelyn Juarez, Emily Thompson, Vaiva Vezys, Melanie E Moses, Judy L. Cannon

## Abstract

T cells are required to clear infection, moving first in lymph nodes to interact with antigen bearing dendritic cells leading to activation. T cells then move to sites of infection to find and clear infection. T cell motion plays a role in how quickly a T cell finds its target, from initial naïve T cell activation by a dendritic cell to interaction with target cells in infected tissue. To better understand how different tissue environments might affect T cell motility, we compared multiple features of T cell motion including speed, persistence, turning angle, directionality, and confinement of motion from T cells moving in multiple tissues using tracks collected with microscopy from murine tissues. We quantitatively analyzed naïve T cell motility within the lymph node and compared motility parameters with activated CD8 T cells moving within the villi of small intestine and lung under different activation conditions.

Our motility analysis found that while the speeds and the overall displacement of T cells vary within all tissues analyzed, T cells in all tissues tended to persist at the same speed, particularly if the previous speed is very slow (less than 2 *µm*/min) or very fast (greater than 8 *µm*/min) with the exception of T cells in the villi for speeds greater than 10 *µm*/min. Interestingly, we found that turning angles of T cells in the lung show a marked population of T cells turning at close to 180*^o^*, while T cells in lymph nodes and villi do not exhibit this “reversing” movement. Additionally, T cells in the lung showed significantly decreased meandering ratios and increased confinement compared to T cells in lymph nodes and villi. The combination of these differences in motility patterns led to a decrease in the total volume scanned by T cells in lung compared to T cells in lymph node and villi. These results suggest that the tissue environment in which T cells move can impact the type of motility and ultimately, the efficiency of T cell search for target cells within specialized tissues such as the lung.

## 1 Introduction

Cell migration is a key feature of cellular function, and T cells are particularly specialized to migrate in different tissue types as infection can occur in any tissue and T cell movement in individual tissues is crucial to clear infection. Prior to infection, naïve T cells move within the paracortex of the lymph node, and upon interaction with cognate antigen bearing dendritic cells, T cells activate and effector CD8 T cells move to peripheral tissue sites of infection. In tissue sites, CD8 T cells enter infected tissues and move within tissues in order to find and kill target cells, including virally infected cells or tumor cells. Interestingly, T cells can move through many tissue environments that differ in cell type, structure, and chemical cues such as chemokines each of which may affect T cell motility patterns. While many studies have identified key molecules that regulate CD8 T cell effector function in different tissues, still relatively little quantitative analysis has been done to analyze the way CD8 T cells navigate multiple different types of tissue environments to find target cells.

CD8 T cell motility is a key feature of CD8 T cell function, particularly in searching through complex tissue environments to identify and interact with target cells. Motility of T cells is a function of a combination of T cell-intrinsic mechanisms, the extracellular environment, and chemical signals in the milieu [31]. Tissue environments include a complex and heterogeneous system of cell types, extracellular matrix components, and soluble factors which have been shown to impact T cell motion; for example, structural cells within the tissue environment provide signals to feed back to immune cells in the central nervous system ([7], [56]). Additionally, multiple studies have shown that naïve T cells in the lymph node paracortex use interactions with fibroblastic reticular cells to mediate movement, as well as receive soluble signals such as IL-7 for survival ([5], [29], [33]). In addition to extracellular influences, many studies have defined intrinsic molecular regulators of T cell movement, particularly speed in multiple tissues including lymph nodes ([21], [29], [23], [15], [13]), skin ([20], [18], [48] [3]), FRT [10], liver ([24], [38], [52]), lung ([42], [2], [16]) just to name a few. High speeds have been linked to integrins (e.g. LFA-1 and VLA-4) [28] [48], chemokine receptors ([4], [45], [3]), as well as signaling molecules such as regulators of the actin cytoskeleton ([42], [43], [12]). These studies identified key molecular drivers and structures that mediate T cell movement within individual tissues, but there remains a gap in analysis to compare how T cell motility patterns might differ between tissues.

Quantitative analysis of cell motion provides a powerful tool to determine underlying mechanisms that drive how cells, including T cells, move. While structures, cell types, and chemical cues may differ in tissues that can impact T cell motility patterns, studies performed both in vitro and in vivo have found that all cell movement, including T cells, use actomyosin contractility and actin flow to couple directional persistence and speed, pointing to a universal mechanism for cells to move faster and more persistently in a direction ([27], [34]). This universal coupling of directional persistence and speed is most clearly shown in cells moving in vitro and on 2D surfaces. How T cells navigate complex tissue environments in three dimensions is still not well understood where cells use multiple modes of migration [62].

In this study, we quantitatively analyze T cell movement as one way to interrogate potential environment influences from different tissues. We previously used quantitative analyses of T cell movement in tissue to reveal specific types of motility patterns leading to more effective T cell responses ([21], [42], [55]). In this paper, we compare multiple features of T cell motion in different tissues: speed, tendency to persist at a speed, dependence of speed on turning angle, mean squared displacement, directionality, confined ratio and time, and volume patrolled within the lymph node with naïve T cells and activated CD8 T cells within the small intestine and lung. By comparing T cell movement in different tissues, we identify tissue specific effects on T cell motility. Our results suggest that tissue environments may contribute to different modes of T cell movement, which can impact the efficiency of T cell searches for target cells in tissues.

## 2 Results

### 2.1 Speed

We began our analysis with a comparison of the cell-based and displacement speeds of T cells in multiple tissues including naïve CD4 and CD8 T cells in the lymph node (LN) in the absence of infection [21] (“LN”), (Video 1); effector CD8 T cells moving in the villi in response to lymphocytic choriomeningitis virus (LCMV) infection at day 8 post infection [55] (“Villi”) (Video 2); effector CD8 T cells moving in LPS inflamed lung at days 7-8 post infection [42] (“Lung LPS”) (Video 3); and effector CD8 T cells in influenza-infected lung at days 7-8 post infection (“Lung Flu”) (Video 4). Specifics about the cell tracks analyzed for each condition is found in Table 1.

**Table 1.** Two-photon microscopy T cell data.

|  | Lymph Node (LN) | Small Intestine (villi) | Lung (Flu) | Lung (LPS) |
| --- | --- | --- | --- | --- |
| Number of T cells | 4400 | 425 | 355 | 191 |
| Type of T cell | Naïve | Activated | Activated | Activated |
| Mode of Activation | Not activated | In vivo | In vitro | In vivo |
| T cell specificity | Polyclonal | TCR transgenic | Polyclonal and TCR transgenic | Polyclonal |
| Imaging modality | Tissue explant | In situ | Tissue explant | In situ and tissue explant |
| Source | Fricke [21] et al. and Tasnim et al. [54] | Thompson et al. [55] | Mrass and Cannon | Mrass [42] et al. |

We previously found no difference in motility speed and patterns of naïve CD4 and CD8 T cells in lymph nodes [21]. To ensure consistency across analyses, we normalized time steps to 90 seconds for all data sets (for details, see Materials and Methods).

Figure 1A and Figure 1B shows the box-and-whisker plot of cell-based speed and displacement speed from each tissue. The whiskers show the *Q*_1_ − 1.5 * (*Q*_3_ − *Q*_1_) and *Q*_3_ + 1.5 * (*Q*_3_ − *Q*_1_) values where *Q*_1_ and *Q*_3_ are the first and third quartiles.

**Fig 1.**
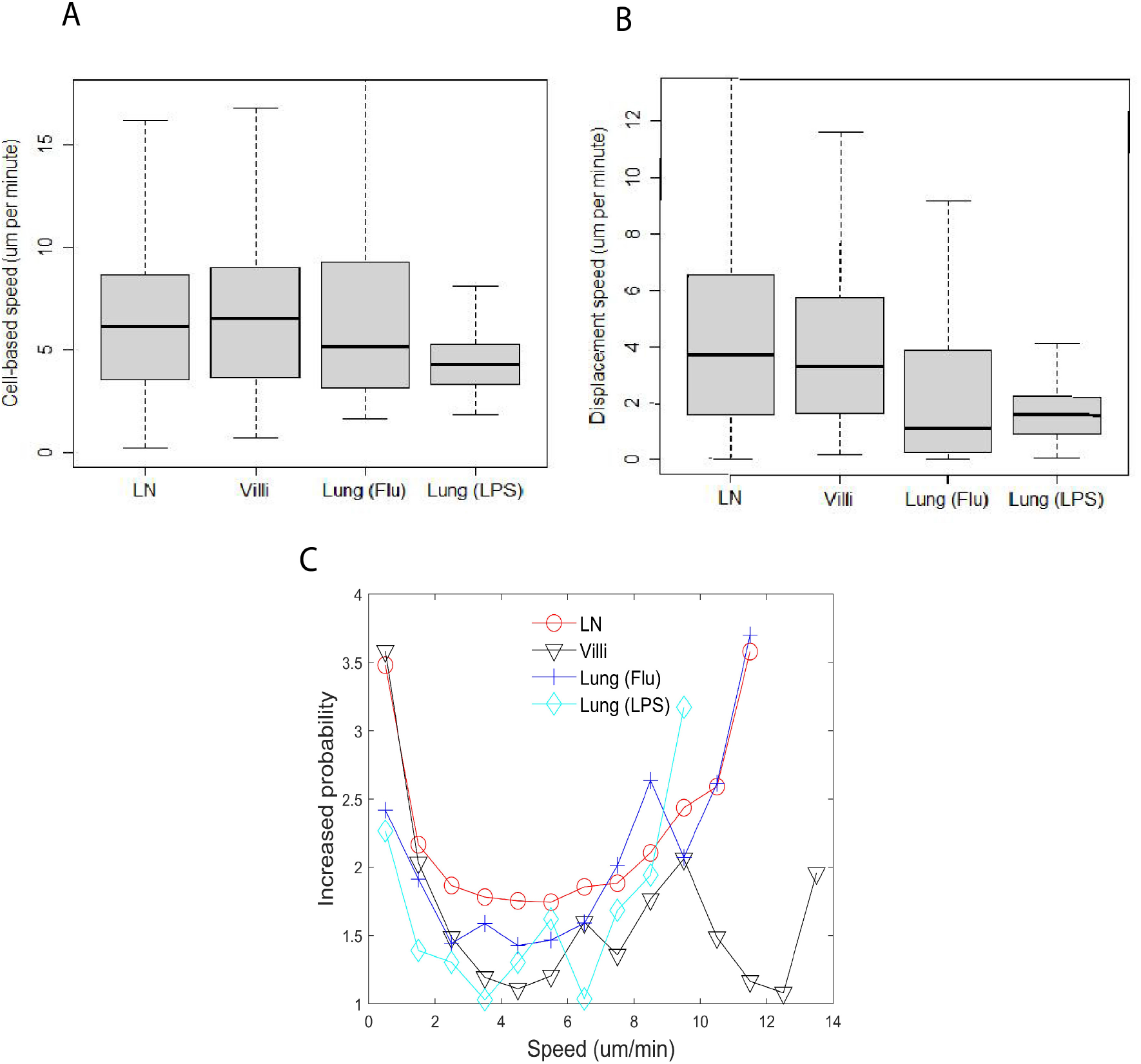
Speed distribution of T cells does not correlate with tissue type or activation status. (A) Box-and-whisker plot of cell-based speed (*µm*/min) of T cells moving in LN (median 6.2), villi (median 6.5), lung (Flu infected) (median 5.2) and lung (LPS) (median 4.3). (B) Box-and-whisker plot of displacement speed (*µm*/min) of T cells in lymph (median 3.7), villi (median 3.3), lung (Flu infected) (median 1.1) and lung (LPS instilled) (median 1.6). (C) Distribution plot of probability to persist at the same speed.

The median cell-based speed for naïve T cells in the LN was 6.2 *µ*m/min, CD8 effector T cells in the villi 6.5 *µ*m/min, CD8 effector T cells from influenza infected lung (Flu) 5.2 *µ*m/min and CD8 effector T cells from LPS-inflamed lung (LPS) 4.3 *µ*m/min. Pairwise p-values based on cell-based average speeds from two different tissues are reported in Table 3. P-values are computed using the paired Wilcoxon Rank Sum test (otherwise known as the Mann-Whitney U test) using the statistical package R with the Bonferroni correction for multiple comparisons. The correction was used for all tables showing p-values.

**Table 2.**
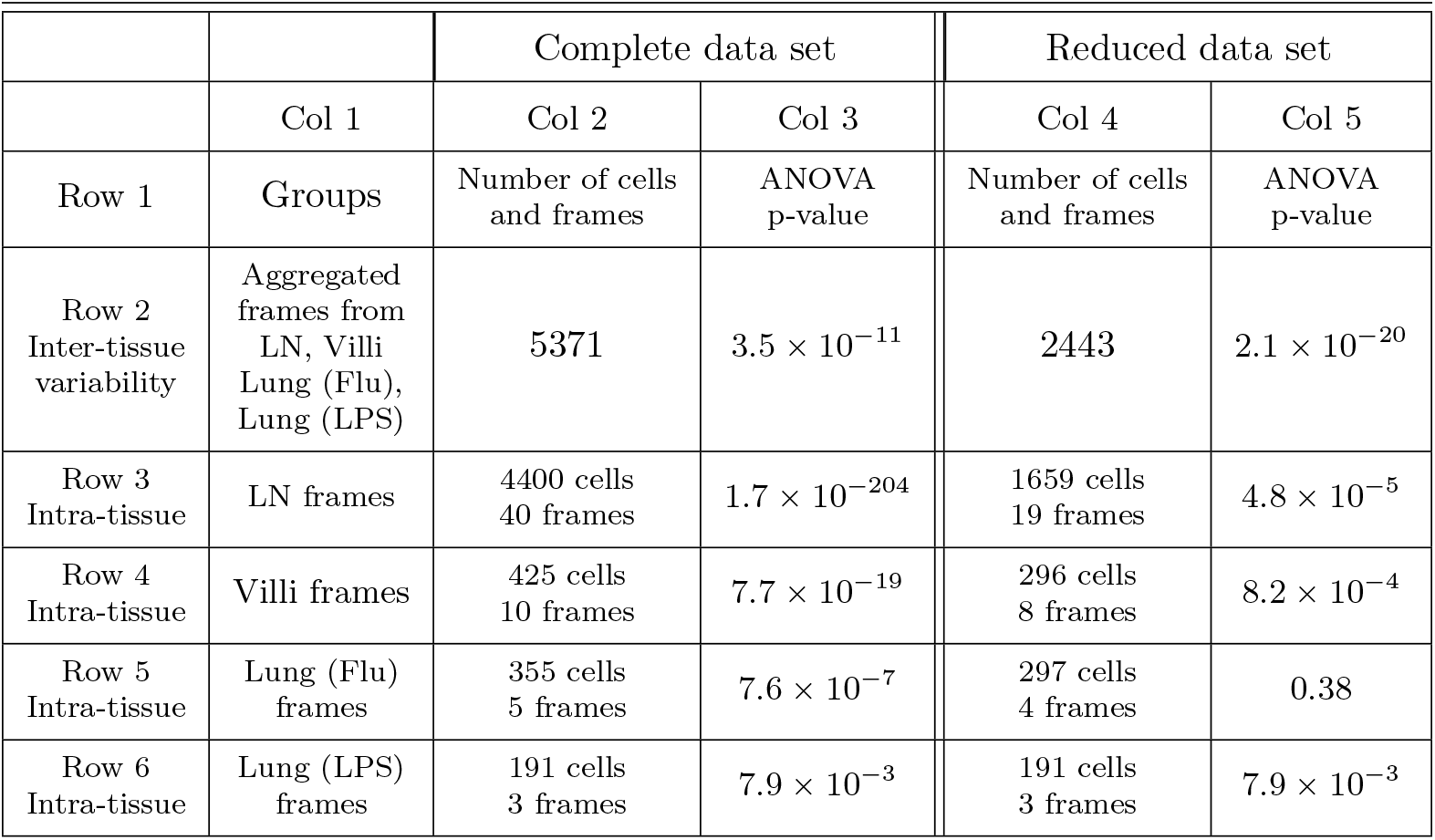
We performed an Analysis of Variance (ANOVA) study which computes a ratio of differences in means between groups and within groups. The ANOVA test uses a F-distribution which computes a ratio of between-groups variance and within-groups variance. We previously showed that in the LN, motility of T cells captured in different fields and on different days from different lymph nodes of different mice did not contribute significantly to variation within T cell movement of the data set [32]. To test intra-tissue variability, the groups consisted of frames composed of two-photon tracks within the same tissue of a single mouse. To test inter-tissue variability, the sets consisted of the aggregated frames of two-photon tracks of all mice imaged in the same tissue. To assess intra-tissue variability compared with inter-tissue variability, we performed an ANOVA analysis of cell-based speed. ANOVA analysis shows that while there exist significant differences in T cell motility between the different tissues (Table 2, Column 3, Row 2), ANOVA analysis also shows that there exist even more significant differences within frames of each individual tissue, particularly of T cells in the lymph node (Table 2, Column 3, Rows 3-6). We found the same trend when performing the ANOVA test with the displacement speed and volume per time. To decrease the variability within a tissue type, we selected a reduced set of frames from each tissue based on statistical variability. Twenty-one frames of the most variable frames were removed from the 40 LN frames which increased the ANOVA p-value from 1.7 × 10^−204^ to 4.8 × 10^−5^ when the ANOVA test was re-run with the remaining nineteen frames using the cell-based speed (Row 3, Column 5). Two variable frames were removed from the ten villi frames which increased the ANOVA p-value from 7.7 × 10^−19^ to 8.2 × 10^−4^ (Row 4, Column 5). One frame was removed from the five Lung (Flu) frames which increased the ANOVA p-value from 7.6 × 10^−7^ to 0.38 (Row 5, Column 5). No frames were removed from the Lung (LPS) data set as none were statistically variable from the other frames. The remaining number of T cells for each tissue is shown in Column 4. The dramatic increase in p-value demonstrates that the removed frames were outliers compared with the data from the same tissue. The data set with outlier frames removed is called “reduced data set”. As the variability within each tissue is reduced, the inter-tissue p-value decreased from 3.5 × 10^−11^ to 2.1 × 10^−20^ in Row 2 when the reduced set of files is analyzed. The new inter-tissue p-value 2.1 × 10^−20^ is significantly smaller than the p-values measuring the intra-tissue variability in Column 5, Rows 3-6 of Table 2.

|  |  | Complete data set |  | Reduced data set |  |
| --- | --- | --- | --- | --- | --- |
|  | Col 1 | Col 2 | Col 3 | Col 4 | Col 5 |
| Row 1 | Groups | Number of cells and frames | ANOVA p-value | Number of cells and frames | ANOVA p-value |
| Row 2<br>Inter-tissue variability | Aggregated frames from LN, Villi Lung (Flu), Lung (LPS) | 5371 | $3.5 \times 10^{-11}$ | 2443 | $2.1 \times 10^{-20}$ |
| Row 3<br>Intra-tissue | LN frames | 4400 cells<br>40 frames | $1.7 \times 10^{-204}$ | 1659 cells<br>19 frames | $4.8 \times 10^{-5}$ |
| Row 4<br>Intra-tissue | Villi frames | 425 cells<br>10 frames | $7.7 \times 10^{-19}$ | 296 cells<br>8 frames | $8.2 \times 10^{-4}$ |
| Row 5<br>Intra-tissue | Lung (Flu) frames | 355 cells<br>5 frames | $7.6 \times 10^{-7}$ | 297 cells<br>4 frames | 0.38 |
| Row 6<br>Intra-tissue | Lung (LPS) frames | 191 cells<br>3 frames | $7.9 \times 10^{-3}$ | 191 cells<br>3 frames | $7.9 \times 10^{-3}$ |

**Table 3.** Table of p-values of pairwise comparisons of cell-based speed as shown in Figure 1A using Wilcoxon rank sum test.

|  | LN | Villi | Lung (Flu) | Lung (LPS) |
| --- | --- | --- | --- | --- |
| LN | 1.0 | 0.51 | 1.0 | $5.4 \times 10^{-14}$ |
| Villi | 0.51 | 1.0 | 1.0 | $7.8 \times 10^{-14}$ |
| Lung (Flu) | 1.0 | 1.0 | 1.0 | $4.9 \times 10^{-5}$ |
| Lung (LPS) | $5.4 \times 10^{-14}$ | $7.8 \times 10^{-14}$ | $4.9 \times 10^{-5}$ | 1.0 |

Figure 1A shows that naïve T cells in lymph nodes (LNs) and effector CD8 T cells in the gut villi moved at similar speeds (Figure 1A; LN: (6.2 *µ*m/min); villi: (6.5 *µ*m/min); Table 3). Both naïve T cells in the LN and effector CD8 T cells in the gut villi moved significantly faster than effector CD8 cells in LPS-inflamed lung (4.3 *µ*m/min) (Table 3). Effector CD8 T cells in the influenza-infected lung moved slightly faster than effector CD8 T cells in the LPS inflamed lung (Figure 1A: lung (Flu) 5.2 *µ*m/min vs lung (LPS) 4.3 *µ*m/min (*p* = 4.9 × 10^−5^). Figure 1 - supplement 1A in the Supplementary Data shows the frequency distribution of cell-based speeds for T cells moving in each tissue type. The figure shows that T cells in either influenza-infected lung or LPS-inflamed lung have a large proportion of cells moving at slower speeds compared to T cells in lymph nodes or villi, with T cells in LPS-inflamed lung showing the largest proportion of cells moving at slow speeds.

We also analyzed effector CD8 T cells moving in the villi at day 5 (d5) post infection and compared with effector T cells moving in villi at day 8 (d8) post infection (Supplementary Data, Figure 1 - supplement 2). A direct comparison of effector T cells moving d5 versus d8 post infection in the villi show that some motility parameters remain similar, including persistence (Figure 1 - supplement 2C). Effector T cells in the villi move slightly faster at d8 compared to d5 (Figure 1 - supplement 2A,B), likely reflecting decreasing antigen load with clearance of virus at later times post infection.

The similar speeds of naïve T cells in LNs and effector T cells in villi at d8 suggests that activation status is not a sole driver of T cell speed in different tissues despite the significant changes in expression of cell surface markers that regulate motility. Naïve T cells express CCR7 and CD62L, while activated T cells upregulate many different cell surface receptors including CD44, CD103, as well as tissue homing chemokine receptors such as CCR9 for gut homing ([35], [36], [46], [60]), and CXCR3 and CXCR4 for lung homing ([49], [30], [17], [59], [39]) . It has been shown that antigen increases interaction time and the difference in speed between effector CD8 T cells on d5 and d8 likely reflect antigen load [25]. However, antigen interaction cannot be the sole driver of speed, as antigen-independent CD8 T cells in the LPS-inflamed lung move more slowly than antigen-specific T cells in all other tissues (Figures 1A and 1B).

Furthermore, the differences in effector T cell speed between T cells moving in LPS-inflamed lung and influenza-infected lung suggest that the specific tissue environment does not fully dictate the speed of T cell movement.

We also analyzed the intra-tissue variation in T cell motility within each tissue using ANOVA (Table 2, Column 3, Rows 3-6 shows the p-values). Our results show that within each tissue, particularly the lymph node, T cells can show high variance in motility. We analyzed motility parameters including cell-based speed when outlier frames with highly variable moving T cells are removed from each tissue (Table 2, Column 5, Rows 3-6). Interestingly, while removing variable frames slightly increases cell-based speed and displacement speed (Figure 1 - supplement 3A,B), the relative differences between tissues is preserved (compare Figure 1 with Figure 1 - supplement 3).

We then calculated displacement speed of T cells in each tissue, which measures the speed at which the cell moves away from an initial location and is smaller than the cell-based speed in all the tissues (Figure 1B). The displacement speed is statistically similar between naïve T cells in LN (median 3.7 *µ*m/min) and effector CD8 T cells within the villi (3.3 *µ*m/min). P-values of comparisons between each T cell type are reported in Table 4. We found that the displacement speed of effector CD8 T cells in the lung in both influenza infection and LPS treatment are similar to each other and both statistically significantly lower than T cells in the LNs and villi (influenza-infected lung 1.1 *µ*m/min, LPS lung 1.6 *µ*m/min). Figure 1 - supplement 1B shows the frequency distribution of T cell displacement speed. These results suggest that the lung environment leads to lower displacement speed of T cells.

**Table 4.** Table of p-values of pairwise comparisons of displacement speed shown in Figure 1B using Wilcoxon rank sum test.

|  | LN | Villi | Lung (Flu) | Lung (LPS) |
| --- | --- | --- | --- | --- |
| LN | 1.0 | 1.0 | $< 2 \times 10^{-16}$ | $< 2 \times 10^{-16}$ |
| Villi | 1.0 | 1.0 | $< 2 \times 10^{-16}$ | $< 2 \times 10^{-16}$ |
| Lung (Flu) | $< 2 \times 10^{-16}$ | $< 2 \times 10^{-16}$ | 1.0 | 0.13 |
| Lung (LPS) | $< 2 \times 10^{-16}$ | $< 2 \times 10^{-16}$ | 0.13 | 1.0 |

### 2.2 Persistence

We calculated the likelihood that an individual T cell will persist in moving at the same speed in each tissue (Figure 1C). We quantified the likelihood that a T cell will continue to move at the same speed as the previous time step and termed this “persistence” (for detailed methods, see Section 4.5). For example, Figure 1C shows that for the T cells in the LN, a T cell is 3.5 times more likely to continue moving at a slow speed (less than 1 *µ*m/min) given that it was moving slowly (less than 1 *µ*m/min) in the previous time step.

We observed a similar trend to persist at very low speeds (< 2 *µm*/min) for T cells moving in all tissues observed and at very high speeds (> 8 *µm*/min) in all the tissues. T cells in the villi exhibited a decrease in persistence at speeds above 10 *µ*m/min while T cells in LN and lung showed similar increase in persistence above 8 *µ*m/min with no decrease at higher speeds. The increased persistence at high speeds in the villi was seen for effector T cells at d5 post infection (Figure 1 - supplement 2C). At very low speeds (< 2 *µm*/min), T cells in the lymph node and villi show a higher likelihood of persistently moving at a slow speed compared to T cells in the lung. Figure 1C shows that at intermediate speeds (between 3 - 7 *µm*/min), T cells in the lung and villi exhibit lower persistence likelihood than T cells in the lymph node, suggesting that the lung and gut environment can hinder the ability of T cells to move persistently at these intermediate speeds.

### 2.3 Mean squared displacement

We then determined the mean squared displacement (MSD) for T cells moving in each tissue. Figure 2A plots the MSD for a representative cell vs time from each tissue type. The linear regression line is shown with the scatter plot whose slope is computed using the log of the MSD and the log of time. We then calculated the slope of the linear regression line for each T cell from a tissue. All T cell slopes from a tissue are used to create the box-and-whisker plot shown in Figure 2B. T cells are tracked for a maximum of 10.5 minutes to ensure consistency of analysis across tissues [31].

**Fig 2.**
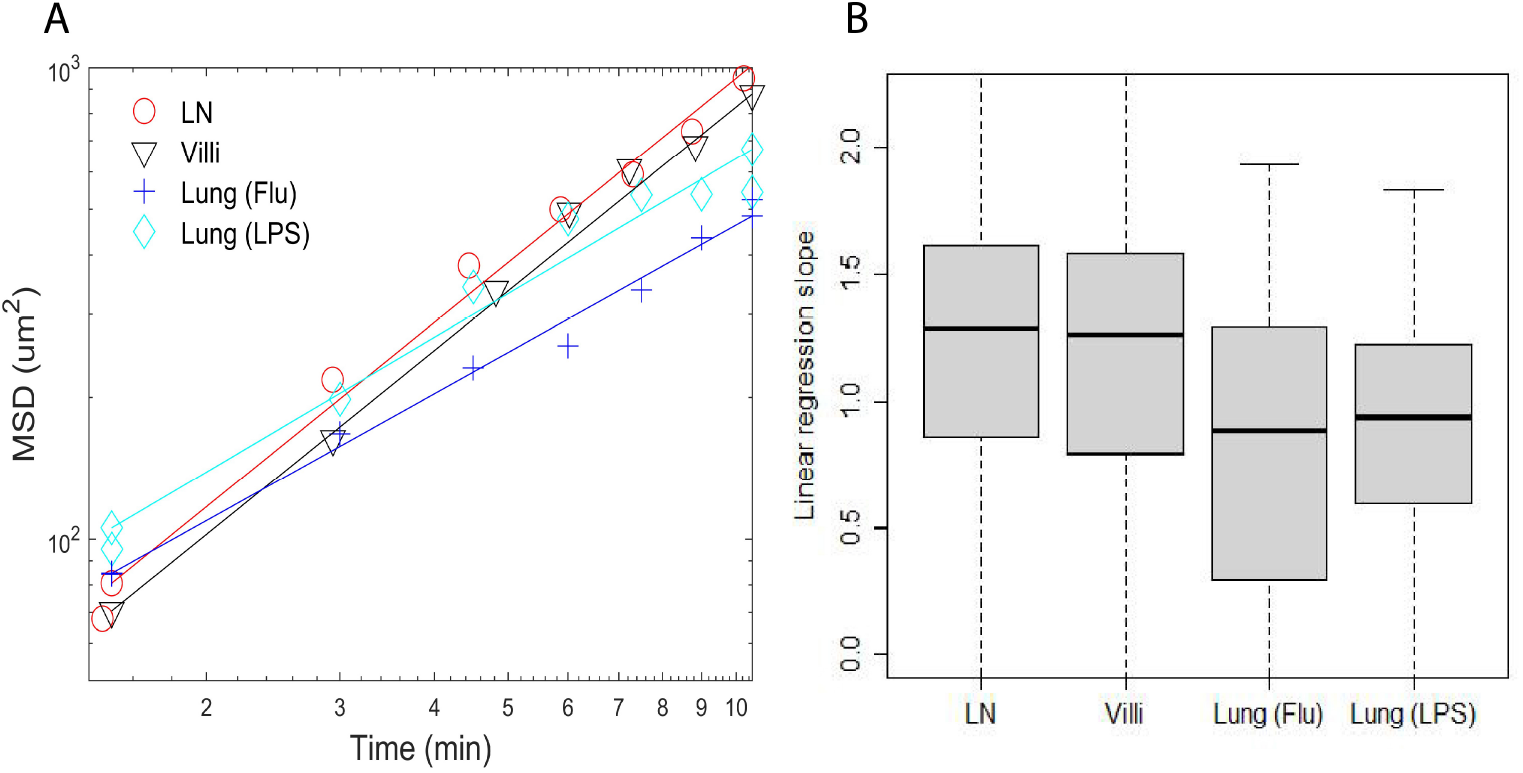
(A) Plots of mean square displacement (MSD) vs time and linear regression lines of individual representative cells near median from Fig. 2B. (B) Box-and-whisker plots of linear regression cell slopes of log transformed mean squared displacement vs time. The median values are LN (1.3), villi (1.3), lung (Flu) (0.88), and lung (LPS) (0.94).

As shown in Figure 2B, T cell motion in the LN and villi could be characterized as superdiffusive with values >1. In contrast, the slope of T cells in the LPS inflamed lung was close to one (0.94) while the slope of T cells in the influenza infected lung is less than one (0.88) and would be considered diffusive and subdiffusive [31]. The p-values comparing the differences between the mean square displacement slopes of T cells moving in individual tissues are shown in Table 5. The slope of MSD between T cells moving in LN and villi were similar, and significantly different from T cells moving in the lung (Table 5). This result remained similar even if outlier frames are removed (Figure 2 - supplement 2A,B).

**Table 5.**
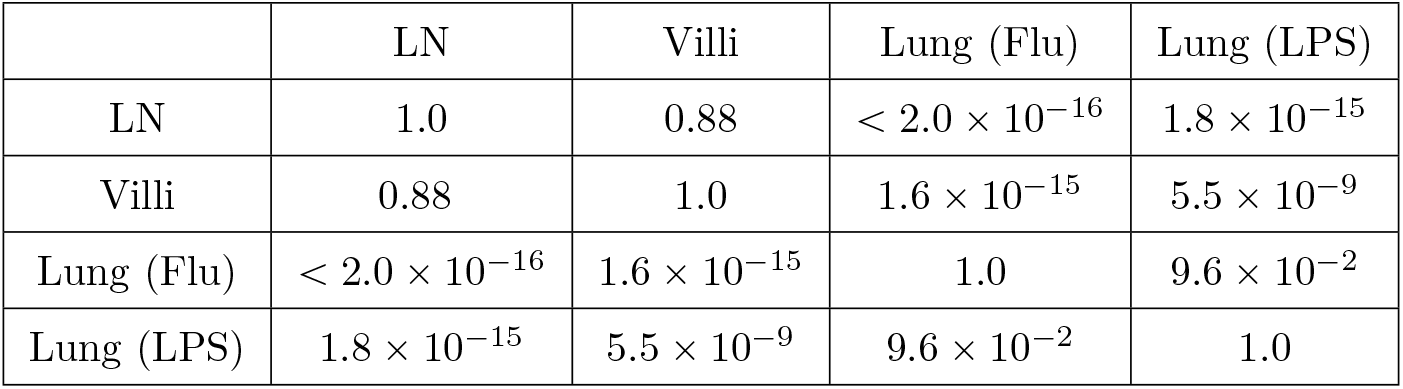
Table showing mean square displacement p-values as pairwise comparisons from Figure 2B using Wilcoxon rank sum test.

### 2.4 Turning angle and dependence of speed on turning angle

As persistence in cell motion is related to turning angles, we analyzed the turning angles of T cells in individual tissues. Figure 3A plots the relative frequency of all turning angles of T cells moving in different tissues. We did not include T cells moving at speeds less than 1 *µm*/min. We reasoned that turning angles are not relevant when a cell is moving very slowly (< 1 *µm*/min). Also small speeds will emphasize turning angles in increments of 45*^o^* degrees due to the pixel resolution of the microscope. Since the distribution is not uniform or flat in Figure 3A, the cell motion cannot be considered Brownian.

**Fig 3.**
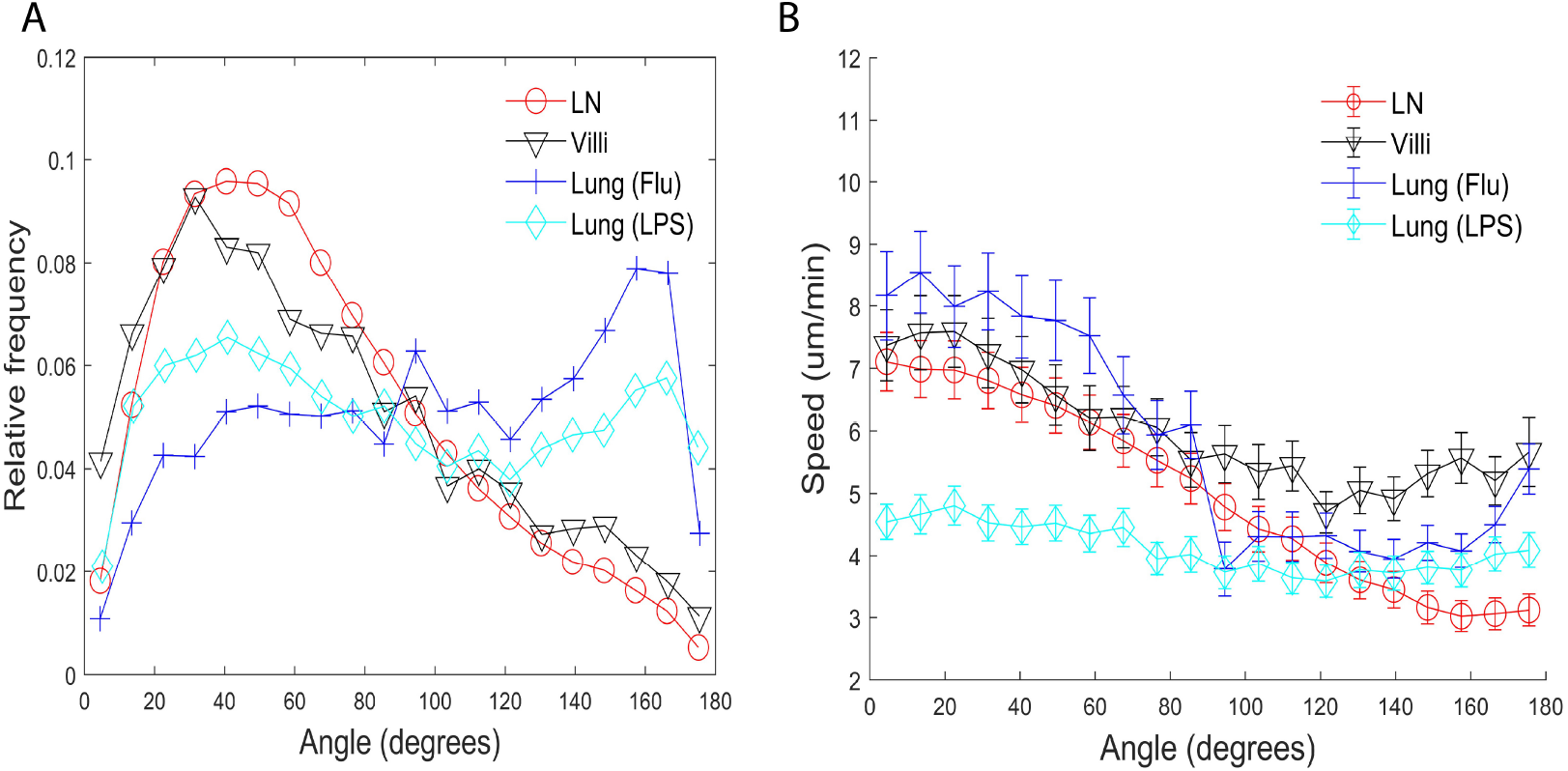
Turning angles and coupling of speed and turning angles of T cells in different tissues. (A) Relative frequency distribution of turning angles in each tissue. T cells moving in the lung show a peak at approximately 160*^o^*. (B) Plot of speed (um/minute) versus angle (degrees). The speed tends to decrease as the turning angle increases in all tissues except for T cells in the LPS-inflamed lung. Error bars show plus and minus 1/8 of the standard deviation within each 9*^o^* angle bin.

We found that while T cells in all tissues show some preference for turning angles between 40*^o^* - 50*^o^*, many more T cells in the lymph node and villi showed the preference to turn at smaller angles compared to T cells in the lung. There was no statistical difference between the LN and villi (See Table 6 for a list of all the p-values.) Interestingly, T cells moving in the lung showed a peak at approximately 160*^o^*, a behavior not seen in T cells moving in lymph node and villi. This peak is likely due to the “back and forth” motion observed in T cells in the lung which we have previously described [42]. The higher percentage of T cells turning at smaller angles in the lymph nodes and villi suggests that these organs allow for a broader range of turning motion, potentially enabling broader search areas.

**Table 6.** Table of p-values of pairwise comparisons of proportion of turning angles less than 90*^o^* shown in Figure 3A using Wilcoxon rank sum test.

|  | LN | Villi | Lung (Flu) | Lung (LPS) |
| --- | --- | --- | --- | --- |
| LN | 1.0 | 1.0 | $< 2.0 \times 10^{-16}$ | $< 2.0 \times 10^{-16}$ |
| Villi | 1.0 | 1.0 | $< 2.0 \times 10^{-16}$ | $< 2.0 \times 10^{-16}$ |
| Lung (Flu) | $< 2.0 \times 10^{-16}$ | $< 2.0 \times 10^{-16}$ | 1.0 | $2.6 \times 10^{-3}$ |
| Lung (LPS) | $< 2.0 \times 10^{-16}$ | $< 2.0 \times 10^{-16}$ | $2.6 \times 10^{-3}$ | 1.0 |

We extended our analysis by determining if there exists a relationship between speed and the turning angle. Maiuri et al. [34] and Jerison and Quake [27] previously found that T cells that move faster generally move persistently in one direction and show a small turning angle while slower T cells show higher turning angles. Our results confirmed that T cells in the lymph node, villi, and influenza-infected lung moving with faster speeds exhibit smaller turning angles while T cells moving with slower speeds exhibit larger turning angles for all tissues (Figure 3B). Interestingly, effector CD8 T cells moving in the LPS-inflamed lung did not show the speed-turning angle correlation (Figure 3B, cyan), suggesting that the relationship between speed and turning angle may not be universal. The behavior in the LPS lung could be due to the fact that T cells in the LPS-inflamed lung have a slow cell-based speed; however, the flatness of the line suggests that even slow T cells in LPS-inflamed lung may not be subject to the same mechanisms that regulate the speed-angle behavior seen in faster moving cells. We also note that T cells in the influenza-infected lung experience a small increase in speeds for turning angles between 160*^o^* and 180*^o^*.

### 2.5 Directionality and Confinement

The turning angle determines the directionality of cell movement. While the displacement speed is lower than the cell-averaged speed in all tissues, the amount of reduction from cell-based to displacement speed differs from tissue to tissue. This suggests that directional persistence in T cell movement may differ in the different tissues analyzed. We assessed directionality by calculating a “meandering ratio” which quantified how likely a T cell deviates from its original direction. Figure 4 shows the box-and-whisker plot of the meandering ratio of T cells moving within the different tissues. T cells in the lymph node and villi move significantly more directionally than T cells in the lung (median values for the meandering ratio are lymph node: 0.70, villi: 0.63, lung (Flu): 0.22, and lung (LPS): 0.37). Pairwise p-value comparisons are reported in Table 7. The meandering ratio remained the same even after outlier frames were removed (Figure 4 - supplement 2).

**Fig 4.**
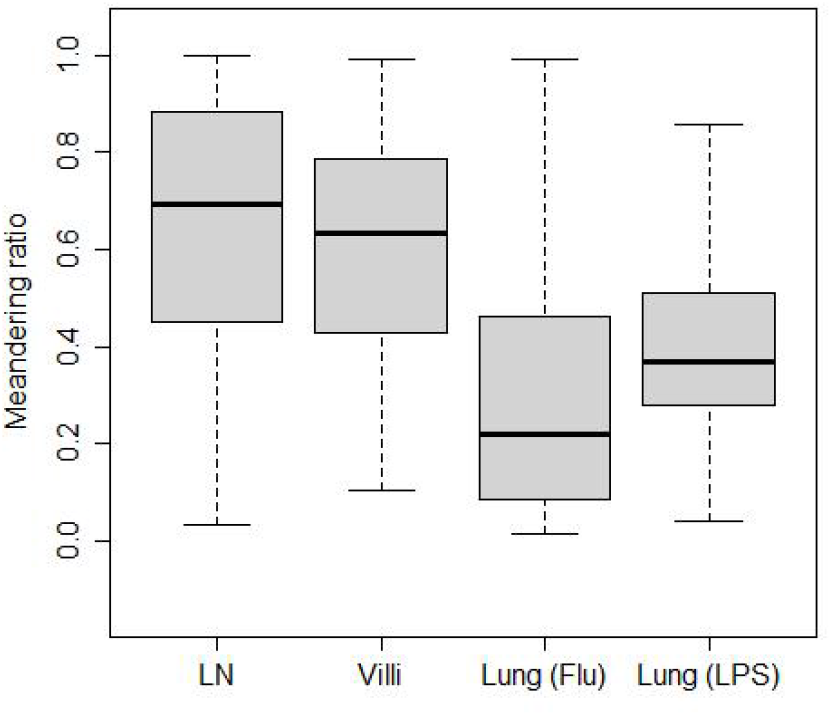
Meandering ratio within different tissues. The median values are LN (0.70), villi (0.63), lung (Flu) (0.22), and lung (LPS) (0.37).

**Table 7.** Table shows p-values of pairwise comparisons of meandering ratio as shown in Figure 4 using Wilcoxon rank sum test.

|  | LN | Villi | Lung (Flu) | Lung (LPS) |
| --- | --- | --- | --- | --- |
| LN | 1.0 | $1.6 \times 10^{-4}$ | $< 2 \times 10^{-16}$ | $< 2 \times 10^{-16}$ |
| Villi | $1.6 \times 10^{-4}$ | 1.0 | $< 2 \times 10^{-16}$ | $< 2 \times 10^{-16}$ |
| Lung (Flu) | $< 2 \times 10^{-16}$ | $< 2 \times 10^{-16}$ | 1.0 | $4.3 \times 10^{-9}$ |
| Lung (LPS) | $< 2 \times 10^{-16}$ | $< 2 \times 10^{-16}$ | $4.3 \times 10^{-9}$ | 1.0 |

We have previously shown that T cells can alternate between confined motion in which the search area is localized and ballistic motion in which motion is fast and persistent in a direction [42]. We calculated the confined ratio as defined by the time a T cell spends confined versus moving in Figure 5A. We found that naïve T cells in the lymph node spend very little time confined and most of the time moving, showing a median confined ratio of 0.15. The confined ratio of effector CD8 T cells in the villi was slightly higher (0.2). Effector CD8 T cells in the influenza infected lung had a confined ratio of 0.53 while effector T cells in LPS-inflamed lung showed the highest confined ratio of 0.60 (p-values are reported in Table 8).

**Fig 5.**
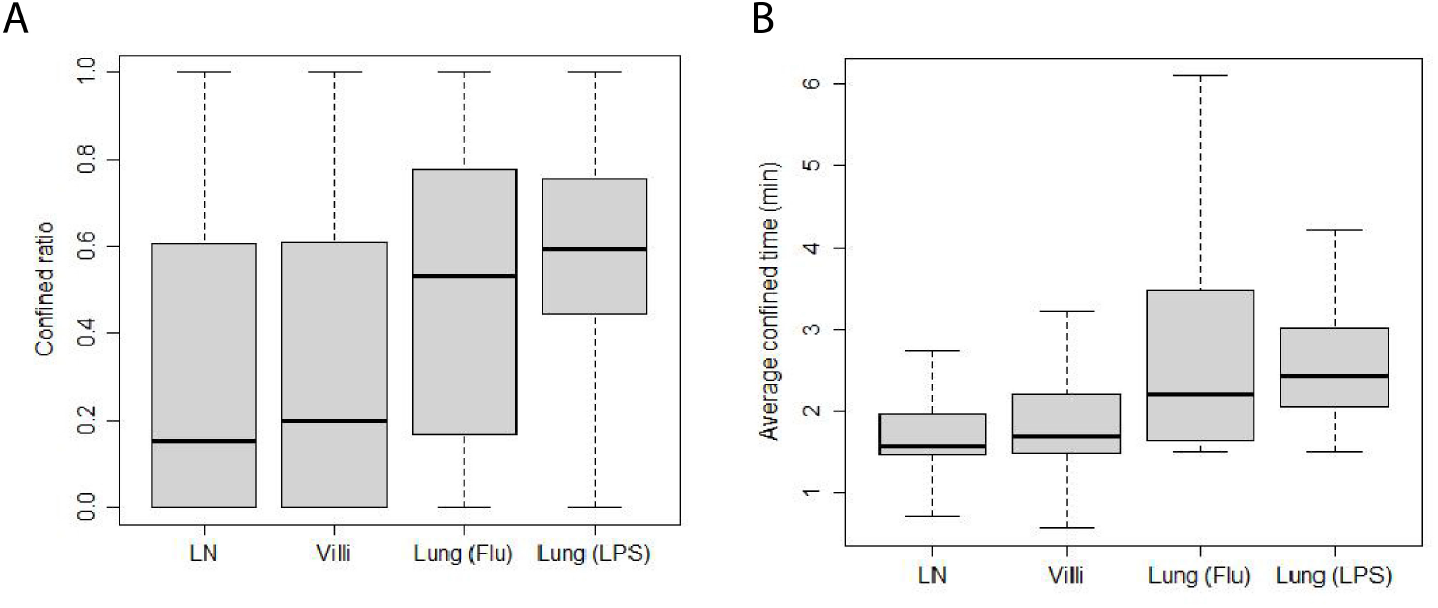
Confinement of T cells from different tissues. (A) Box-and-whisker plot of confined ratios. Median values: LN 0.15, villi 0.2, lung (Flu) 0.53, lung (LPS) 0.60. (B) Box-and-whisker plot of confined time. Median values (minutes): LN 1.6, villi 1.7, lung (Flu) 2.2, lung (LPS) 2.4.

**Table 8.** Table showing p-values of pairwise comparisons of confined ratios from Figure 5A using Wilcoxon rank sum test.

|  | LN | Villi | Lung (Flu) | Lung (LPS) |
| --- | --- | --- | --- | --- |
| LN | 1.0 | 1.0 | $< 2 \times 10^{-16}$ | $< 2 \times 10^{-16}$ |
| Villi | 1.0 | 1.0 | $1.3 \times 10^{-11}$ | $< 2 \times 10^{-16}$ |
| Lung (Flu) | $< 2 \times 10^{-16}$ | $1.3 \times 10^{-11}$ | 1.0 | $1.8 \times 10^{-2}$ |
| Lung (LPS) | $< 2 \times 10^{-16}$ | $< 2 \times 10^{-16}$ | $1.8 \times 10^{-2}$ | 1.0 |

We also calculated the average amount of time T cells from each tissue spend confined which is reported as confined time in Figure 5B (p-values reported in Table 9). We found that T cells in the lung (Flu) and lung (LPS) showed significantly longer confined times than T cells in lymph node or villi. Effector T cells in the villi at d5 post infection showed significantly higher confinement ratio and confined time compared with d8 (Figure 5 - supplement 1A,B). Both confined ratios and confined time remained similar even if outlier frames were removed (Figure 5 - supplement 2A,B).

**Table 9.**
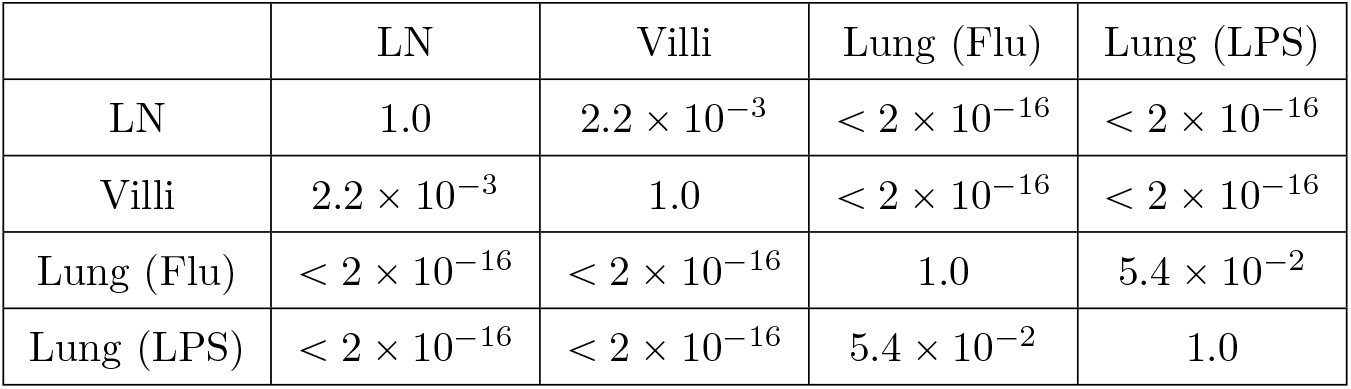
Table showing p-values of pairwise comparisons of confined time from Figure 5B using Wilcoxon rank sum test.

These data show that effector CD8 T cells in LPS-inflamed lung were most confined followed by T cells in influenza-infected lung (Flu) and T cells in the villi. Naïve T cells in the lymph node were the least confined. The low confinement of naïve T cells in the lymph node environment may contribute to the high meandering ratio while confinement as well as antigen in the lung (flu) likely decrease the ability of T cells to move directionally (Figure 4) and lead to low meandering ratios.

### 2.6 Patrolled volume per time

A key function of T cell movement is surveillance of tissues. To assess whether differences we identified in cell speed, directionality, turning angle, and confined ratio ultimately translate into differences in the ability of T cells to survey tissue, we calculated the volume per time patrolled by a T cell residing in different tissues.

Volume per time is a way of incorporating all the different motility parameters we previously identified. Figure 6A shows the amount of volume per time patrolled by the T cells in individual tissues. The volume surveyed is highest for naïve T cells in lymph node (median 9.4 *µm*^3^/s) and d8 CD8 effector T cells in villi (9.4 *µm*^3^/s). Effector CD8 T cells in villi at d5 were intermediate at 6.5 *µm*^3^/s (Figure 6 - supplement 1A). Effector CD8 T cells in the lung showed the lowest volume patrolled: the volume patrolled by T cells in the influenza-infected lung (5.3 *µm*^3^/s) was statistically similar to the volume patrolled by T cells in the LPS-inflamed lung (5.1 *µm*^3^/s). See Table 10 for the p-values.

**Fig 6.**
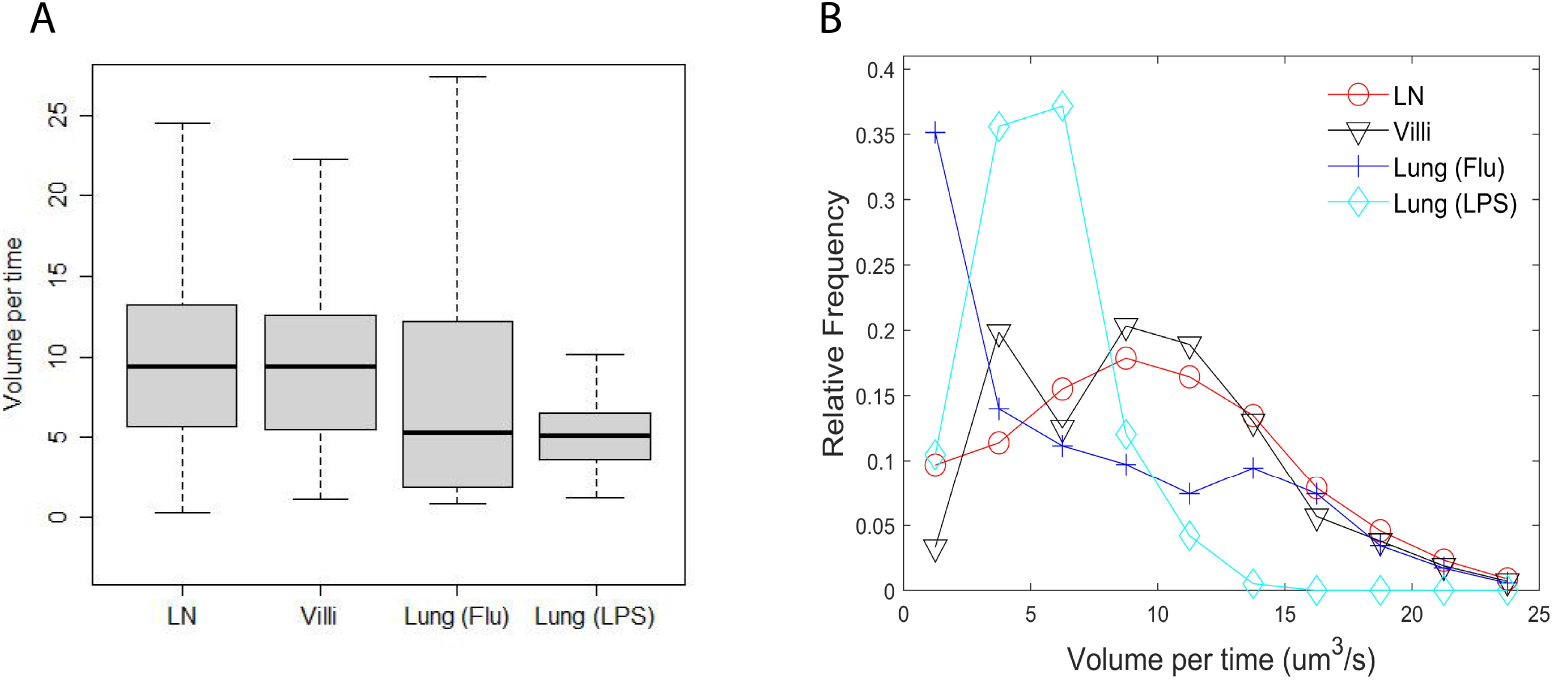
Volume patrolled by T cells in different tissues. (A) Box-and-whisker plot of median volume per time (cubic microns per second) patrolled by T cells in LN (9.4), villi (9.4), lung (flu) (5.3), and lung (LPS) (5.1). (B) Relative frequency distribution of volume per time (cubic microns per second) patrolled by T cells in each tissue.

**Table 10.** Table of p-values of pairwise comparisons of volume per time from Figure 6A using Wilcoxon rank sum test.

|  | LN | Villi | Lung (Flu) | Lung (LPS) |
| --- | --- | --- | --- | --- |
| LN | 1.0 | 1.0 | $< 2 \times 10^{-16}$ | $< 2 \times 10^{-16}$ |
| Villi | 1.0 | 1.0 | $6.9 \times 10^{-12}$ | $< 2 \times 10^{-16}$ |
| Lung (Flu) | $< 2 \times 10^{-16}$ | $6.9 \times 10^{-12}$ | 1.0 | 1.0 |
| Lung (LPS) | $< 2 \times 10^{-16}$ | $< 2 \times 10^{-16}$ | 1.0 | 1.0 |

We also analyzed the full distribution of volume patrolled by T cells in each individual tissue (Figure 6B). The full distribution showed that T cells in lymph nodes and villi at d8 post infection show the largest volume patrolled per time, with T cells in villi at d5 post infection showing a similar distribution of volume scanned (Figure 6 - supplement 1B). Interestingly, although the median patrolled volume is similar between T cells in the LPS-inflamed lung (LPS) and T cells in influenza-infected lung (flu), T cells in the influenza infected lung actually show a large number of cells patrolling at both low and large volumes while T cells in LPS-inflamed lung mostly show low patrol volumes. The Kolmogorov-Smirnov test compares the distributions and shows statistically significant differences in all pairwise comparisons (Table 11). The results were similar with outlier frames removed (Figure 6 -supplement 2A,B).

**Table 11.** Table of p-values of pairwise comparisons of volume per time distributions from Figure 6B using Kolmogorov-Smirnov test.

|  | LN | Villi | Lung (Flu) | Lung (LPS) |
| --- | --- | --- | --- | --- |
| LN | 1.0 | 0.07 | $< 2.2 \times 10^{-16}$ | $< 2.2 \times 10^{-16}$ |
| Villi | 0.07 | 1.0 | $< 2.2 \times 10^{-16}$ | $< 2.2 \times 10^{-16}$ |
| Lung (Flu) | $< 2.2 \times 10^{-16}$ | $< 2.2 \times 10^{-16}$ | 1.0 | $3.7 \times 10^{-9}$ |
| Lung (LPS) | $< 2.2 \times 10^{-16}$ | $< 2.2 \times 10^{-16}$ | $3.7 \times 10^{-9}$ | 1.0 |

These data demonstrate that the combination of speed, turning angle, directional movement, and confinement times all contribute to the ability of T cells to search tissue environments for potential targets.

## 3 Discussion

Cell movement through tissue is an important feature in immunity, particularly for T cell mediated immunity as T cells must make direct cell-cell contact with target cells. A distinguishing feature of T cells is their ability to move through many different tissue environments, which include varying cell types, structural features, and chemokines, all of which may contribute to differences in T cell motility. T cell activation also leads to significant changes in expression of cell surface proteins as well as cytoskeletal machinery that impact T cell motility. Naïve T cells express CCR7 and CD62L ([19], [4], [61]). Upon activation, effector T cells upregulate different chemokine receptors depending on the tissue the effector T cells homes to as well as integrins and CD44 ([49], [30], [17], [59], [39], [35], [46] [36], [60], [55]). Despite these differences, we can use motility parameters to compare the types of movement T cells take independently of the specific molecular interactions that drive these movement patterns.

We analyzed motility parameters to determine what key factors drive the ability of T cells to move through tissues to effectively mount immune responses. Our analysis more fully captures key features of movement that enable T cells to effectively move in tissues, promoting T cell responses. Our study included naïve CD8 and CD4 T cells in the lymph node, antigen specific effector CD8 T cells responding to infection in the villi and lung (influenza-infected lung), and non-antigen specifically activated effector CD8 T cells in the LPS-inflamed lung in a model of acute lung injury [42].

We identified similarities and differences that contribute to the ability of T cells to search tissue for target cells. Interestingly, T cells tend to exhibit similar persistence trends in a range of speeds in all tissue types regardless of activation status. In regards to cell-based speed, non-antigen specific naïve T cells in lymph nodes and antigen specific effector CD8 T cells in villi move fastest while effector CD8 T cells that are non-antigen specific move slowest in the LPS-inflamed lung. These data suggest that the speed at which T cells move can be independent of antigen specific interactions as well as activation status. These results are supported by previous work showing that effector CD8 T cells in the skin appear to move slowly [3] while effector CD8 T cells in the female reproductive tract move at speeds similar to naïve T cells in lymph nodes [10].

Previous work quantitating cell motility in both non-T cells as well as T cells observed a “universal coupling” between speed and directional persistence, showing that fast moving cells show directional persistence while slow moving cells do not ([34], [27]). In our analysis, we find that T cells moving in lymph node, small intestine villi, and influenza-infected lung all show this coupling. However, we have also identified an exception to this rule for T cells moving in the LPS-inflamed lung, which showed no change in directional persistence as measured by turning angle between fast moving and slow moving cells (Figure 3B). Thus, while our data confirm that while speed can be coupled to directional persistence for T cells moving in all tissues analyzed, this coupling is not necessarily “universal”. Recently, it has been shown that the correlation between speed and turning angle can arise from differences in sampling rates [22]. Our data is unlikely to be affected by sampling rate as we equalized and normalized the sampling rate for all the T cells (see Materials and Methods).

We find that T cells moving in the lung show specific motility features that differ from T cells moving in lymph nodes or villi. T cells in the lung tend to displace less, turn at higher angles, particularly at angles > 140*^o^*, meander more, and linger at locations longer. Confinement can occur in the lung independent of antigen, as CD8 T cells activated in vitro in the LPS-inflamed lung can still show confinement without specific antigen activation. In particular, T cells in the lung exhibit back and forth motion, with a peak in turning angles occurring near 160*^o^*. This peak in turning angle is consistent with the “back and forth” motion we previously observed for T cells in the LPS-inflamed lung, as well as a stop-and-go motion [42]. Stop-and-go behavior has also been observed albeit for shorter time periods for T cells in lymph nodes by Miller et al. [41], Wei et al. [58], and Beltman et al. [8]. We also find that the slope of the MSD vs time for T cells moving in lymph nodes and villi show super-diffusive motion as previously observed for effector T cells in the brain [26]. In contrast, the slope of T cells moving in lung is slightly less than 1, suggesting Brownian type motion. However, because the angle distribution is not uniform, the cell motion cannot be considered strictly Brownian. Together these motility parameters suggest that the lung environment may lead to specific types of motion taken by T cells, potentially due to the particular physical environment of the lung.

Our results also show that there can be significant variability of T cell movement patterns within an individual tissue (Table 2). However, despite large differences amongst individual T cell movements within any specific tissue, the overall patterns of T cell motility remain similar within tissue and across tissues.

The volume patrolled by a T cell is dependent not only on its speed but also on turning angles, the cell’s tendency to meander, and the amount of time the cell spends confined to a location. The higher percentage of T cells turning at smaller angles in the lymph nodes and villi suggests that these organs allow a broader range of turning motion, potentially enabling broader search areas as reflected in the larger volumes patrolled. We found that naïve T cells and CD8 effector cells in the villi at d8 post infection patrol a larger volume (9.4 *um*^3^/sec) due to a combination of fast speed, less confinement, more super-diffusive motion, and smaller turning angles. Analysis of effector T cells at days 5 and 8 post infection in the gut suggest that antigen abundance likely increases confinement and thus decrease T cell speed, leading to a smaller volume patrolled at d5 post infection despite comparable values in meandering ratio and MSD. The patrol volume is significantly smaller for effector T cells in the lung due to greater confinement, more Brownian-like motion, and a greater proportion of large turning angles, particularly angles that suggest a “back and forth” motion.

However, T cells in the influenza-infected lung and LPS-inflamed lung show differences in the frequency distribution of volume per time, with T cells in the influenza-infected lung showing more T cells patrolling at higher volumes than T cells in LPS-inflamed lung. These results suggest that a combination of back and forth motion and slightly faster cell-based speeds can affect the search efficiency for T cells in the influenza-infected lung. Previous results using computational modeling suggests that intermittent and back and forth motion can improve search times [9]. Effector T cells moving in LPS-infected lung show the lowest volume covered (5.1 *um*^3^/s) due to low speeds, higher turning angles, low directionality and high confinement times. The lack of coupling between speed and turning angle may also lead to low search efficiency.

Three-dimensional migration of T cells is a complex interplay of internal cell signaling, the surrounding extracellular tissue environment, molecular signaling and chemokines. We have quantitatively analyzed how T cells move in different tissues using multiple metrics. These metrics provide a way of quantitatively capturing underlying complex features of three-dimensional T cell movement.

## 4 Materials and Methods

Table 1 summarizes the number of T cells tracked within the lymph node (LN), small intestine villi, and lung which were analyzed. Cell tracks were obtained from at least 2 separate fields from at least 2 independent experiments. The data comes in the form of x-, y-, and z-coordinates for each T cell at different time frames.

The lymph node T cell tracks were obtained from data in [21] and [54]. Video 1 in the Supplementary Data shows a representative movie from one field from data analyzed in [21]. The T cell tracking in the small intestine was previously described in Thompson et al. [55]. Video 2 reproduces day 8 (d8) T cell movement in villi from Video S1 shown in [55]. T cells from LPS inflamed lung were previously described in Mrass et al. [42]. Video 3 shows CD8 T cells moving in LPS-inflamed lung from [42].

For T cells from influenza infected lung, mice were infected intranasally with 1 × 10^3^ PFU HKx31 (Charles River). To ensure sedimentation of the virus into the lower respiratory tract, infection was performed while mice were under anesthesia with 90 mg/kg ketamine and 8.1mg/kg xylazine. In some experiments mice received polyclonal naïve GFP+CD8+ T cells from Ubiquitin-GFP animals before infection with influenza. GFP+CD8+ naive T cells were derived from single cell suspensions isolated from spleen and lymph nodes of Ubiquitin GFP animals, and CD8+ T cells isolated using the CD8a+ T Cell Isolation Kit (Miltenyi Biotec), then transferred via the tail vein into recipient mice. Recipient mice received approximately 10^4^ GFP+CD8+ T cells. All work was done in accordance with approved protocols per IACUC institutional approvals.

For imaging of GFP+CD8+ T cells in influenza infected lungs, mice were euthanized and lungs from influenza infected mice were removed at days 7 or 8 post infection. After opening the chest cavity, the lungs were inflated with 2% low melting agarose (Sigma-Aldrich, A0701) at a temperature of 37 degrees. We injected one ml of the solution via a catheter through an incision of the trachea. After inflation, the opening in the trachea was sealed with a knot and agarose solidification was induced by exposing the lungs to a PBS solution with a temperature of 4 degrees Celsius.

After harvest, the lungs were transferred into an incubator and transferred within a biosafety cabinet into a POC-R imaging chamber (LaCon). Imaging was performed with a Zeiss LSM800 Airyscan Confocal Microscope. Due to the transparency of the prepared lung tissue, it was possible to visualize at tissue depths of more than 60 µm. Video 4 shows GFP+CD8+ T cells moving at d8 post infection in influenza-infected lung. In some experiments, imaging was performed with similar setups using a Prairie Ultima Two-photon microscope or a Zeiss LSM 510 microscope. We captured equivalent T cell behavior with the different microscope setups.

Below we summarize the time step sampling and the various metrics used to analyze the T cell motion. These metrics include speed (cell-based and displacement speed), tendency to persist at the same speed, mean squared displacement, directionality through the meandering ratio, confined ratio and time, and volume patrolled per time.

### 4.1 Sampling

Due to the different time steps used in the two-photon microscopy of the different tissues, we sample the position data every 90 seconds (or as close to 90 seconds as possible) for each of the tissues to normalize and equalize T cell analyses. Also for tissues which were sampled at a higher frequency, we are able to use all the data for results involving the turning angle by revisiting times that are skipped in an initial 90 second sampling. For example, suppose observations are made at the following times {*t*_0_*, t*_1_*, t*_2_*, t*_3_*, t*_4_*, …*} where *t*_0_ = 0 *s*, *t*_1_ = 45 *s*, *t*_2_ = 90 *s*, *t*_3_ = 135 *s*, *t*_4_ = 180 *s* and so on. The first sampling of the data retains {*t*_0_*, t*_2_*, t*_4_*, …*} and the second sampling uses {*t*_1_*, t*_3_*, t*_5_*, …*}. We do not sub-sample the LPS-inflamed lung data since the time steps are initially 90 seconds. After this sampling is done, the lymph node mean time step was 89.9 seconds with a standard deviation of 2.9 seconds, the villi mean time step was 93.0 seconds with a standard deviation of 7.9 seconds, the lung (Flu) mean time step was 90.0 seconds with a standard deviation of 0.12 seconds, and the unsampled lung (LPS) mean time step was 90.0 seconds with a standard deviation of 0.10 seconds.

### 4.2 Speed

If (*x_i_, y_i_, z_i_*) refers to the position of a cell at time *t_i_* and (*x*_*i*+1_, *y*_*i*+1_, *z*_*i*+1_) refers to the position of the cell at time *t_i+1_*, let *d*_*i,i*+1_ represent the distance between the two positions

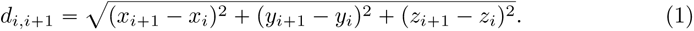

Two different types of speeds are computed using the positions and times from a T cell: a cell-based speed and displacement-based speed.

#### 4.2.1 Cell-based speed

If a cell is tracked for *n* positions and times, the cell-based speed *s_cell_* of a cell is computed by summing all the distances traveled by the cell and dividing by the total elapsed time *t_n_*- *t*_1_,

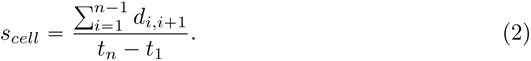

#### 4.2.2 Displacement speed

The displacement speed *s_displacement_* is computed using the first and last locations of the cell,

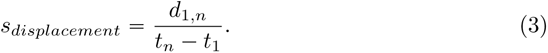

### 4.3 Turning angle and directionality

The turning angle *θ* for a T cell given the three positions of the cell {1,2,3} enclosed within circles is shown in Figure 7A. If **v**_1_ is the vector formed from positions (*x*_1_*, y*_1_*, z*_1_) and (*x*_2_*, y*_2_*, z*_2_), **v**_1_ = (*x*_2_ − *x*_1_*, y*_2_ − *y*_1_*, z*_2_ − *z*_1_) and **v**_2_ is the vector formed from positions (*x*_2_*, y*_2_*, z*_2_) and (*x*_3_*, y*_3_*, z*_3_), **v**_2_ = (*x*_3_ − *x*_2_*, y*_3_ − *y*_2_*, z*_3_ − *z*_2_), the turning angle *θ* is computed using

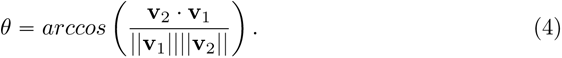

**Fig 7.**
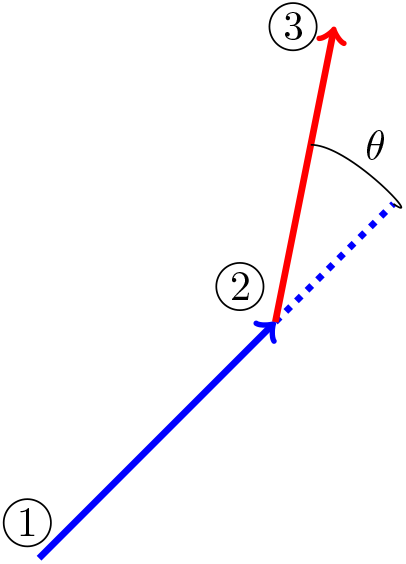
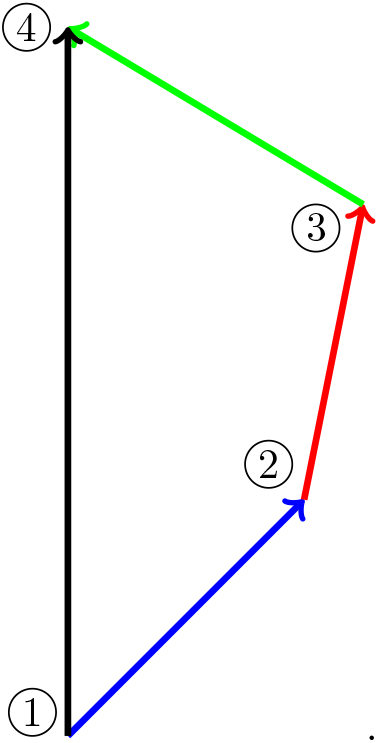
**a.** Turning angle illustration **b.** Meandering ratio illustration

We did not include speeds less than 1 *µm*/min as these cells are likely to be considered stopped and turning angles with very small speeds can lead to artifactual angle measurements; for example, small speeds will emphasize turning angles in increments of 45*°* degrees due to the pixel resolution of the microscope.

Suppose a T cell visits the four locations enclosed within the circles (Figure 7B). One can compute the total distance travelled by summing up the distance from location 1 to location 2, *d*_1,2_, the distance from location 2 to location 3, *d*_2,3_, and the distance from location 3 to location 4, *d*_3,4_. The straight line distance can also be computed from the original location 1 to the final location 4, *d*_1,4_. One measure of a cell’s tendency to maintain its direction is the meandering ratio [16]

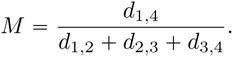

If the ratio *M* is close to 1, the cell deviates very little from one direction, whereas if *M* is much less than 1, the cell moves along a meandering path. In general for *n* locations, directionality can be measured using the ratio

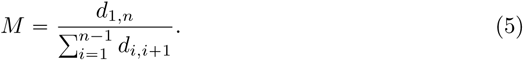

### 4.4 Confined ratio and confined time

We denote the amount of time a T cell lingers in one location as confined time. Given a time *t_i_* and location (*x_i_, y_i_, z_i_*), we count the time difference between *t_i_*and *t_j_* > *t_i_*as confined time if the difference *t_j_* − *t_i_* is greater than 150 seconds and the distance *d_i,j_*is less then 5 *µm*. Once the cell exits (say at time *t_k_*) the 5 *µm* radius centered about (*x_i_, y_i_, z_i_*), the cell is tracked anew and the confined time is computed from *t_k_* and location (*x_k_, y_k_, z_k_*). We call the ratio of confined time to the total time the confined ratio.

In regards to confined time, we calculate the amount of time required to leave a 5 *µm* radius for each cell position. The time is then averaged over all positions to find the confined time.

### 4.5 Tendency to persist at a speed

Let *s_b_* and *s_a_* be two consecutive frame speeds (before and after) in *µm* per minute from a T cell track. If *B_i_* represents the event that *s_b_* lies between *i µm* per minute and (*i* + 1) *µm* per minute, then the probability of event *B_i_* occurring is *P* (*B_i_*) = *m_b_*/*m*, where *m_b_* represents the number of times *i µm*/min ⩽ *s_b_* < (*i* + 1) *µm*/min and *m* represents the total number of tracks. If *A_i_*represents the event that *i µm*/min ⩽ *s_a_*< (*i* + 1) *µm*/min, then the probability *P* (*A_i_*) = *m_a_*/*m* where *m_a_* represents the number of times *i µm*/min ⩽ *s_a_* < (*i* + 1) *µm*/min. Finally it follows that *P* (*A_i_ and B_i_*) = *m_ab_*/*m* where *m_ab_* represents the number of times both criteria are satisfied: *i µm*/*min ⩽ s_a_, s_b_* < (*i* + 1) *µm*/*min* in consecutive frames. According to the definition of conditional probability *P* (*A_i_*|*B_i_*) = *P* (*A_i_ and B_i_*)/*P* (*B_i_*) = *m_ab_*/*m_b_*. The increased likelihood of persisting at the same speed is then calculated as the ratio *P* (*A_i_*|*B_i_*)/*P* (*A_i_*) = (*m* . *m_ab_*)/(*m_a_* . *m_b_*).

### 4.6 Mean squared displacement

Values of the log of the mean squared displacement (*d*_1_*_,n_*)^2^ (MSD) are plotted against the log of the elapsed time. We limit the elapsed time to 10.5 minutes. The slope of the linear regression line is computed from the scatter plot and used to characterize the type of motion. Slope values near 1.0 are associated with Brownian motion, values between 1.0 and 2.0 are associated with Lévy walks, and values less than 1.0 are considered subdiffusive [31].

### 4.7 Rate of volume patrolled

The volume patrolled by a T cell is computed by dividing a 400 *µm* × 400 *µm* × 400 *µm* volume within which a T cell moves into 2.5 *µm* × 2.5 *µm* × 2.5 *µm* cubes. If the distance between a cube center and the T cell center is less than 5 *µm*, the cube volume is assumed to be patrolled. We also connect each two successive cell positions with a straight line and assume the cell patrols volume along the straight line. The total volume patrolled is then divided by the time the T cell is tracked.

### 4.8 Statistical methods

When comparing two groups, p-values were computed using the paired Wilcoxon Rank Sum test (otherwise known as the Mann-Whitney U test) using the statistical package R with the Bonferroni correction for multiple comparisons.

ANOVA was used to assess both intra-tissue and inter-tissue variability. ANOVA uses the F distribution which computes a ratio of variability between groups to variability within groups and is commonly used to test differences between more than two groups. The F value increases as the means between groups increases and the variability within the groups decreases. The larger the F value, the smaller the p-value. To test intra-tissue variability, the groups consisted of frames composed of two-photon tracks within the same tissue of a single mouse. To test inter-tissue variability, the groups consisted of the aggregated frames of two-photon tracks of all mice imaged in the same tissue. These results are discussed in more detail in Table 2. We show that while significant variability does exist within the frames of a tissue, the variability can be reduced be eliminating outliers. Our results using the reduced set of files is similar to the complete set of files (for details, see Supplementary Data Figure 1 - supplement 3, Figures 2-6 supplement 2, and the associated tables within the figures).

## Supporting information

Data format

Lymph node positions

Lung (flu) positions

Lung (LPS) positions

Villi Day 5 positions

Villi Day 8 positions

## 5 Funding

This work was supported by an Institutional Development Award (IDeA) from the National Institute of General Medical Sciences of the National Institutes of Health under Grant P20GM103451 (DJT); NIH 1R01AI097202 (JLC), the Spatiotemporal Modeling Center (P50GM085273), the Center for Evolution and Theoretical Immunology 5P20GM103452 (JLC), AIM CoBRE at UNM HSC NIH P20GM121176 (JLC; PM), NIH 5 T32 AI007538-19 (JRB), institutional support from dedicated health research funds from UNM SOM (JLC), Women in Science Award from UNM, DARPA/AFRL FA8650-18-C-6898 (JLC) and in part institutional support from Dedicated Health Research Funds from the UNM School of Medicine (JLC and PM); and support by the University of New Mexico Comprehensive Cancer Center Support Grant NCI P30CA118100 (JLC and PM).

## 6 Material, Code Availability, and Ethics Statement

All data is available upon request to the corresponding or senior author. All analyzed data is also uploaded to: https://www.biorxiv.org/content/10.1101/2022.11.17.516891v2.supplementary-material. The code used to analyze the T cell tracks can be found at: https://github.com/davytorres/T-cell-analysis-tool. The manuscript includes experiments using animals and all procedures were approved by the IACUC committee, IACUC Animal approval #: 21-201165-HS.

## Supplementary Data

**Figure 1 - supplement 1:**
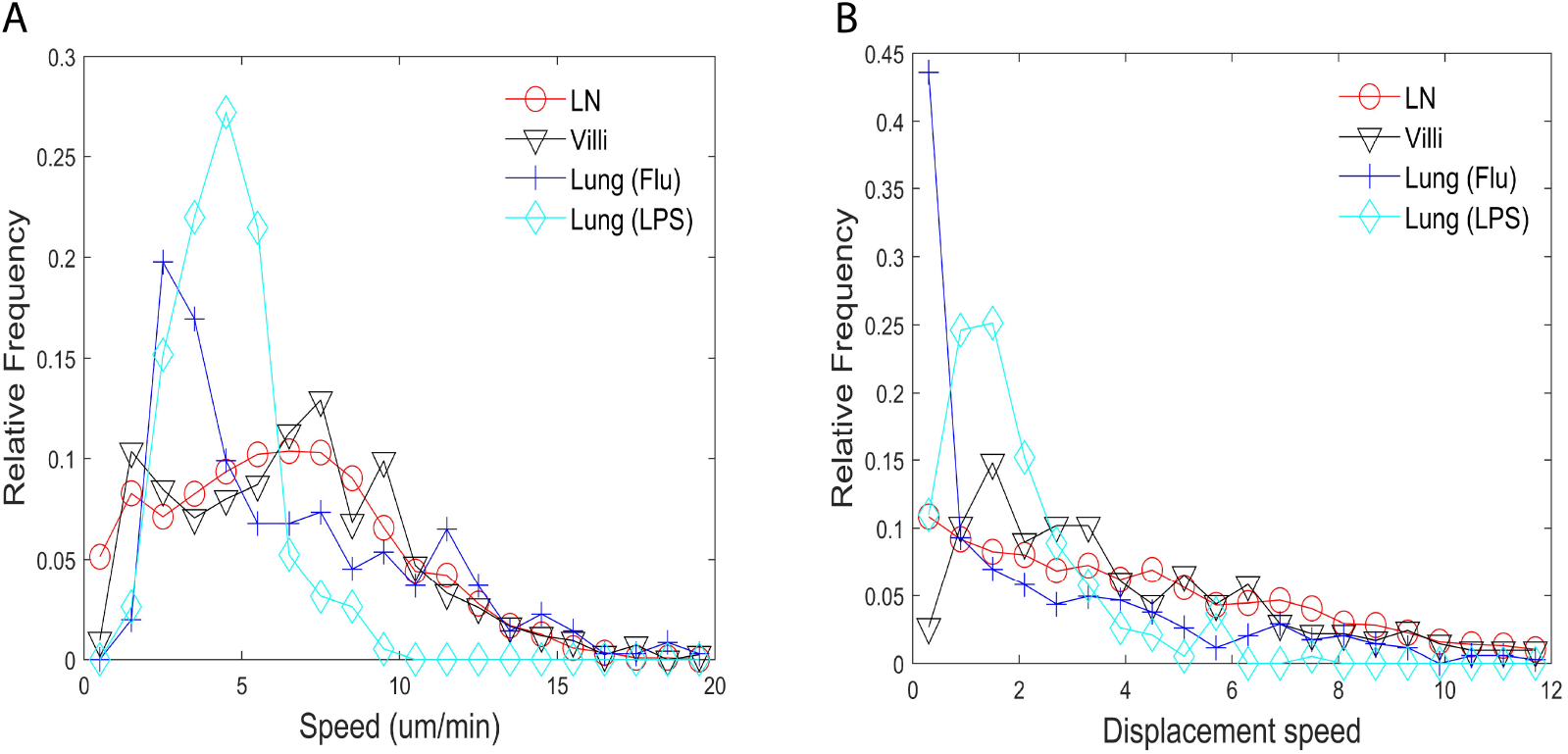
Speed distribution of cell-based (A) and displacement (B) distributions. (A) Relative frequency distribution of cell-based speed in lymph, villi, lung (Flu activated) and lung (LPS activated). The distributions use the average cell-based speed of each T cell. (B) Relative frequency distribution of T cell displacement speed in lymph, villi, lung (Flu activated) and lung (LPS activated).

**Figure 1 - supplement 2:**
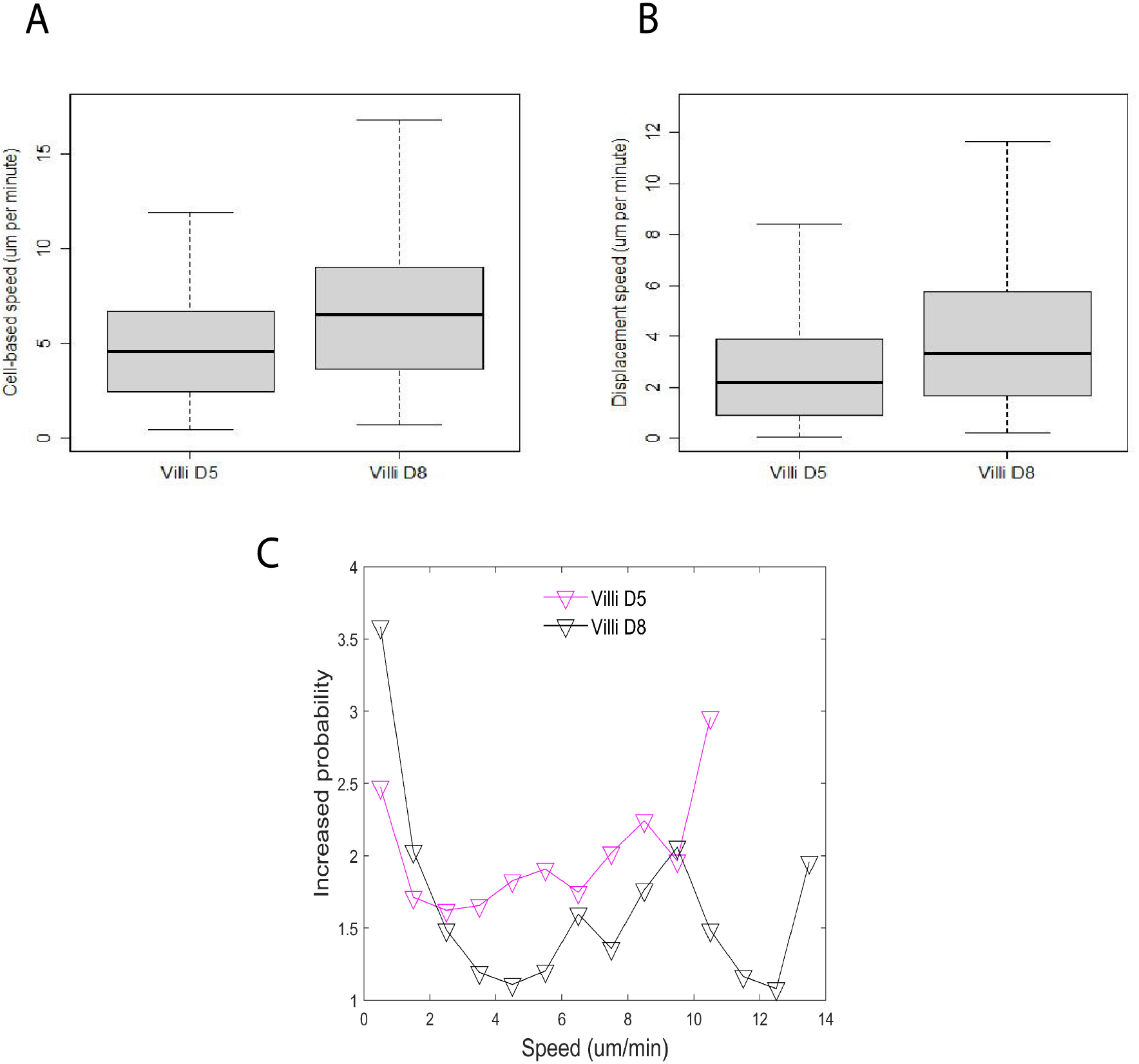
(A) Box-and-whisker plot of cell-based speed (*µm*/min) of T cells moving in villi d5 (median 4.6), villi d8 (median 6.5), p-value < 2.0 × 10^−16^. (B) Box-and-whisker plot of displacement speed (*µm*/min) of T cells in d5 villi (median 2.2), d8 villi (median 3.3), p-value < 2.0 × 10^−16^. (C) Distribution plot of probability to persist at the same speed.

**Figure 2 - supplement 1:**
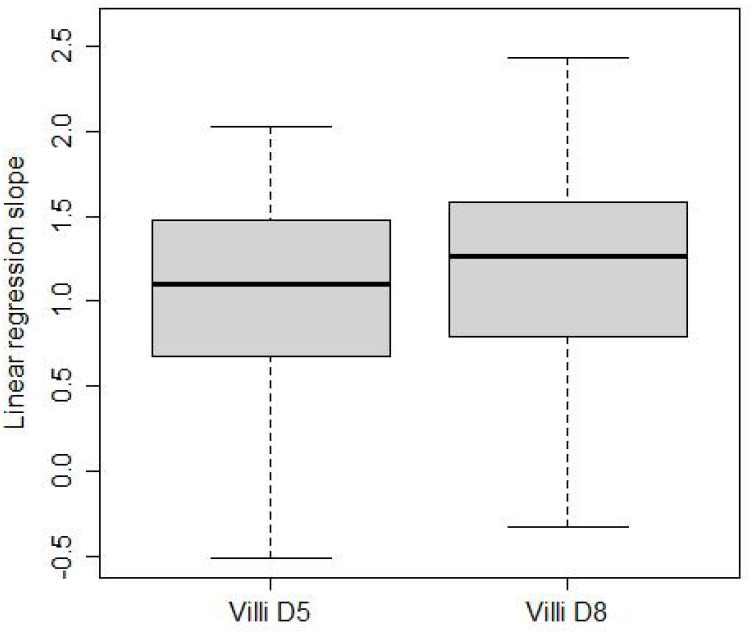
Mean squared displacement linear regression slope is computed for T cells in the villi at d5 and d8 post infection. Box-and-whisker plots of linear regression cell slopes of log transformed mean squared displacement vs time. The median values are villi d5 (1.1), villi d8 (1.3), p-value = 2.6 × 10^−4^.

**Figure 3 - supplement 1:**
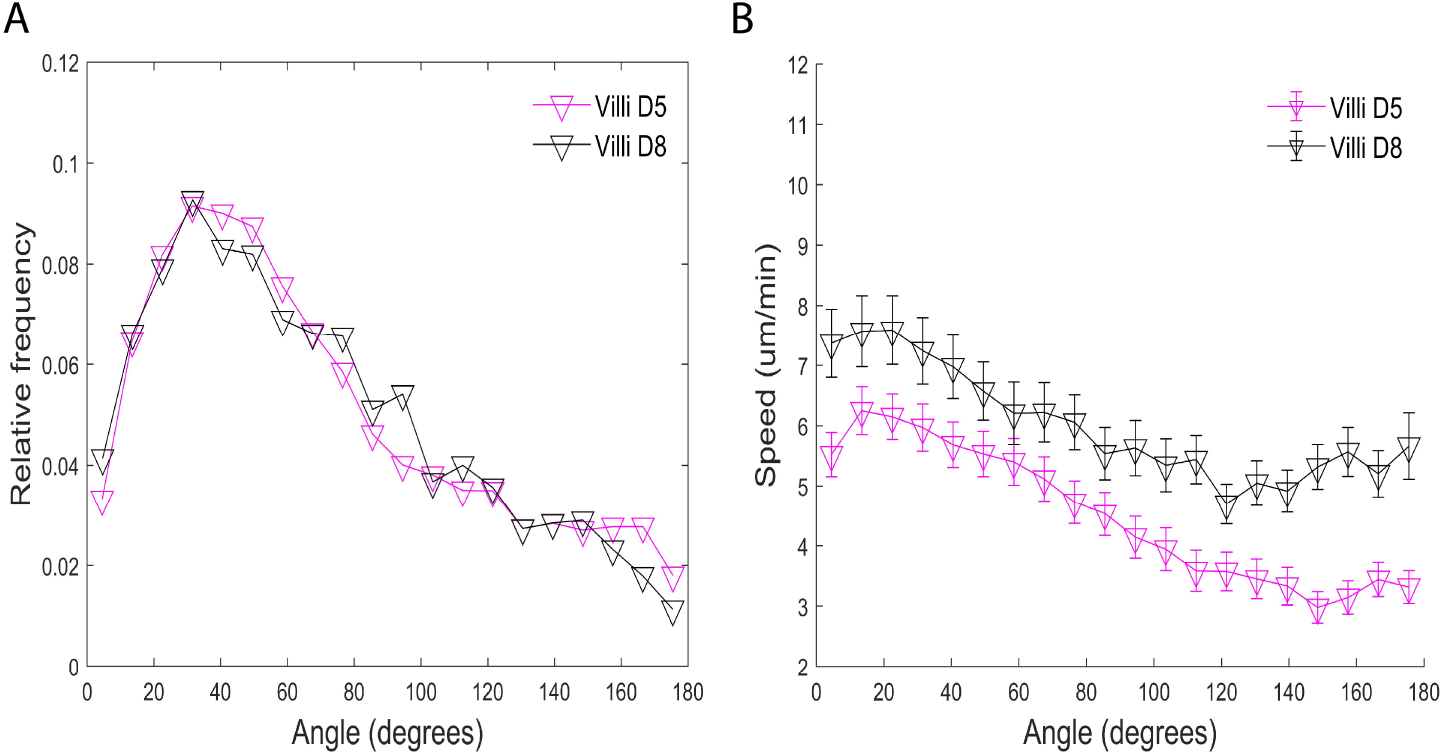
Turning angles and coupling of speed and turning angles of d5 versus d8 CD8 T cells in villi post infection. (A) Relative frequency distribution of turning angles in each tissue. (B) Plot of speed (um/minute) versus angle (degrees). The speed tends to decrease as the turning angle increases. Error bars show plus and minus 1/8 of the standard deviation within each 9*^o^* angle bin.

**Figure 4 - supplement 1:**
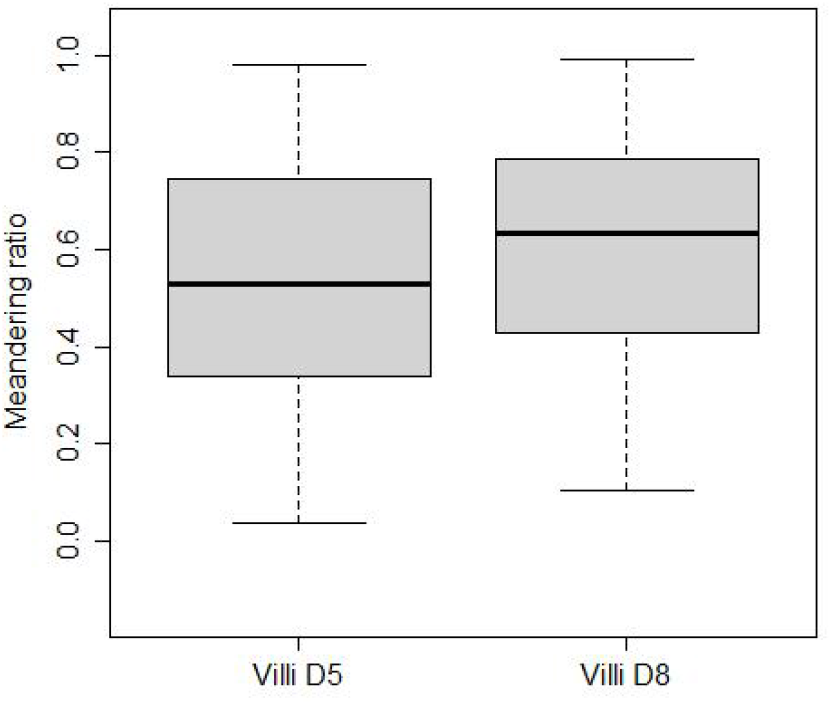
Meandering ratio of CD8 T cells within villi of d5 versus d8 post infection. The median values are villi d5 (0.53), villi d8 (0.63), p-value = 1.1 × 10^−7^.

**Figure 5 - supplement 1:**
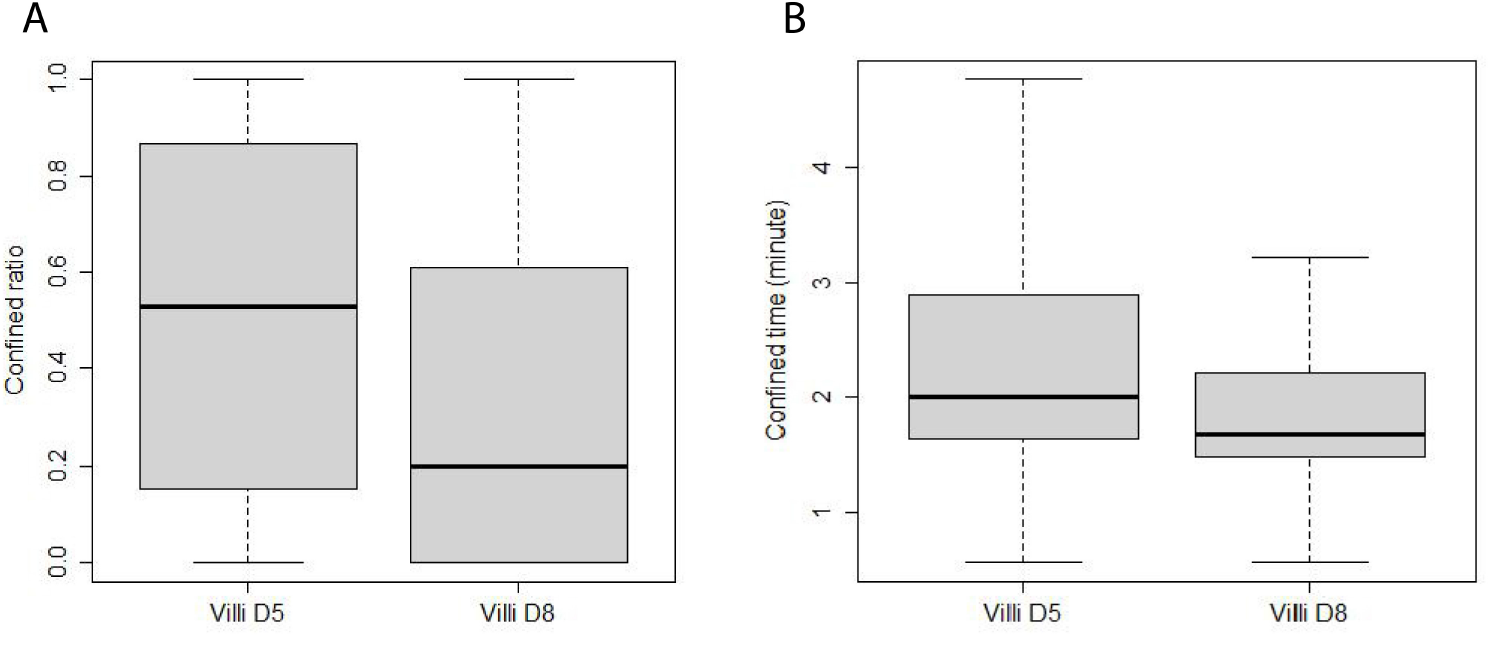
Confinement of T cells from villi d5 versus d8 post infection. (A) Box-and-whisker plot of confined ratios. Median values: d5 villi (0.53), d8 villi (0.2), p-value < 2.0 × 10^−16^ . (B) Box-and-whisker plot of confined time. Median values (minutes): d5 villi (2.0), d8 villi (1.7), p-value < 2.0 × 10^−16^.

**Figure 6 - supplement 1:**
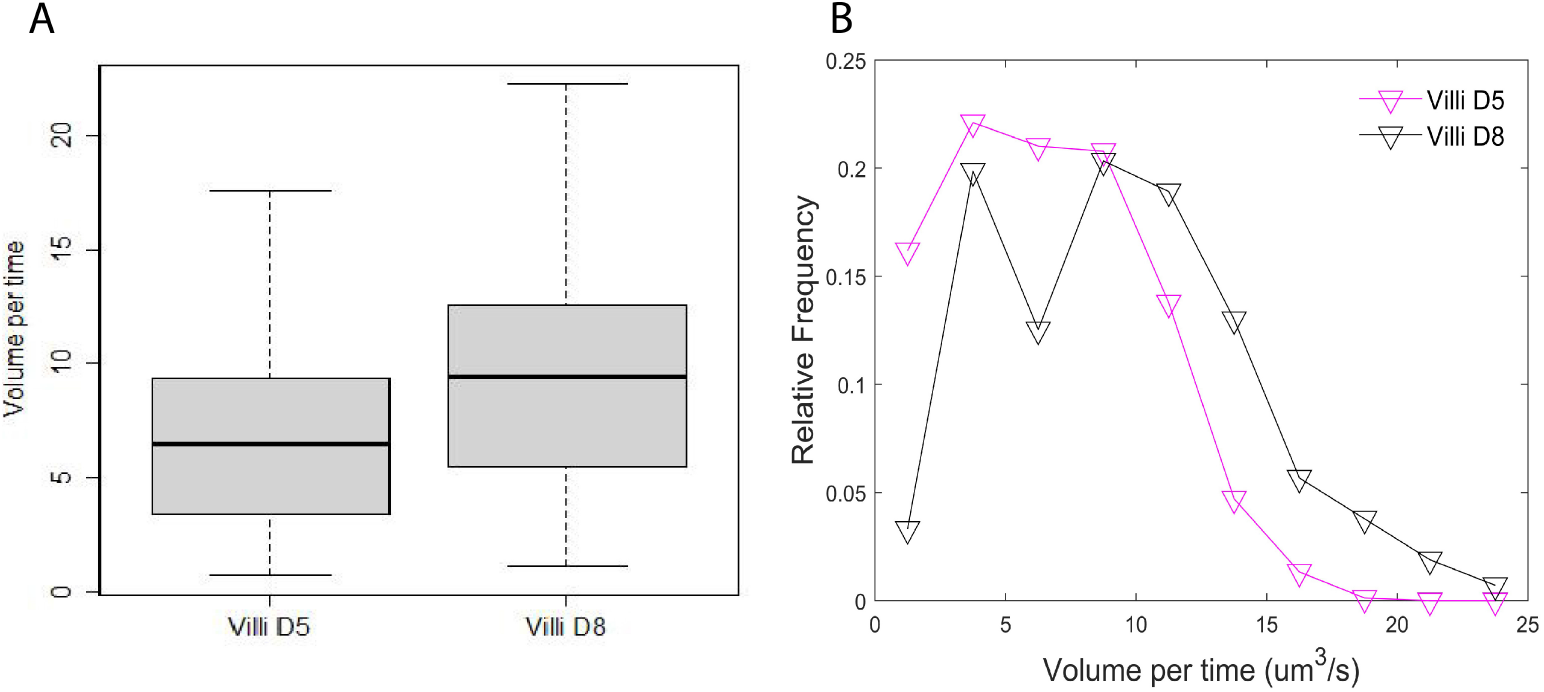
Volume patrolled by d5 versus d8 T cells in villi post infection. (A) Box-and-whisker plot of median volume per time (cubic microns per second) patrolled by T cells in villi d5 (6.5), villi d8 (9.4), p-value < 2.0 × 10^−16^. (B) Relative frequency distribution of volume per time (cubic microns per second) patrolled.

**Figure 1 - supplement 3:**
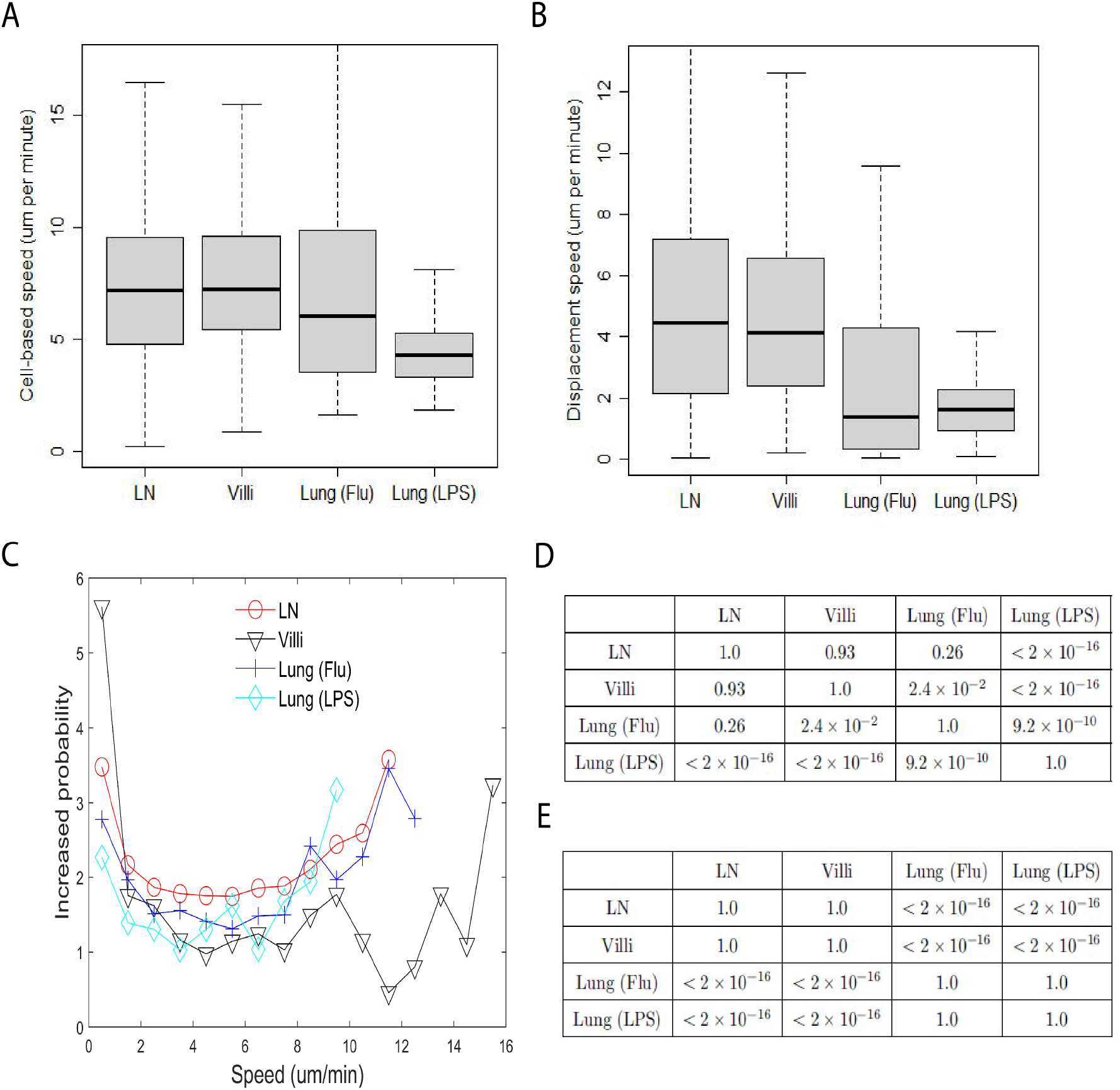
Reduced data set (A) Box-and-whisker plot of cell-based speed (*µm*/min) of T cells moving in LN (median 7.2), villi (median 7.2), lung (Flu infected) (median 6.0) and lung (LPS) (median 4.3). (B) Box-and-whisker plot of displacement speed (*µm*/min) of T cells in lymph (median 4.5), villi (median 4.1), lung (Flu infected) (median 1.4) and lung (LPS instilled) (median 1.6). (C) Distribution plot of probability to persist at the same speed. (D) Table of p-values of pairwise comparisons of cell-based speed as shown in Figure 1A - supplement 3 using Wilcoxon rank sum test. (E) Table of p-values of pairwise comparisons of displacement speed shown in Figure 1B - supplement 3 using Wilcoxon rank sum test.

**Figure 2 - supplement 2:**
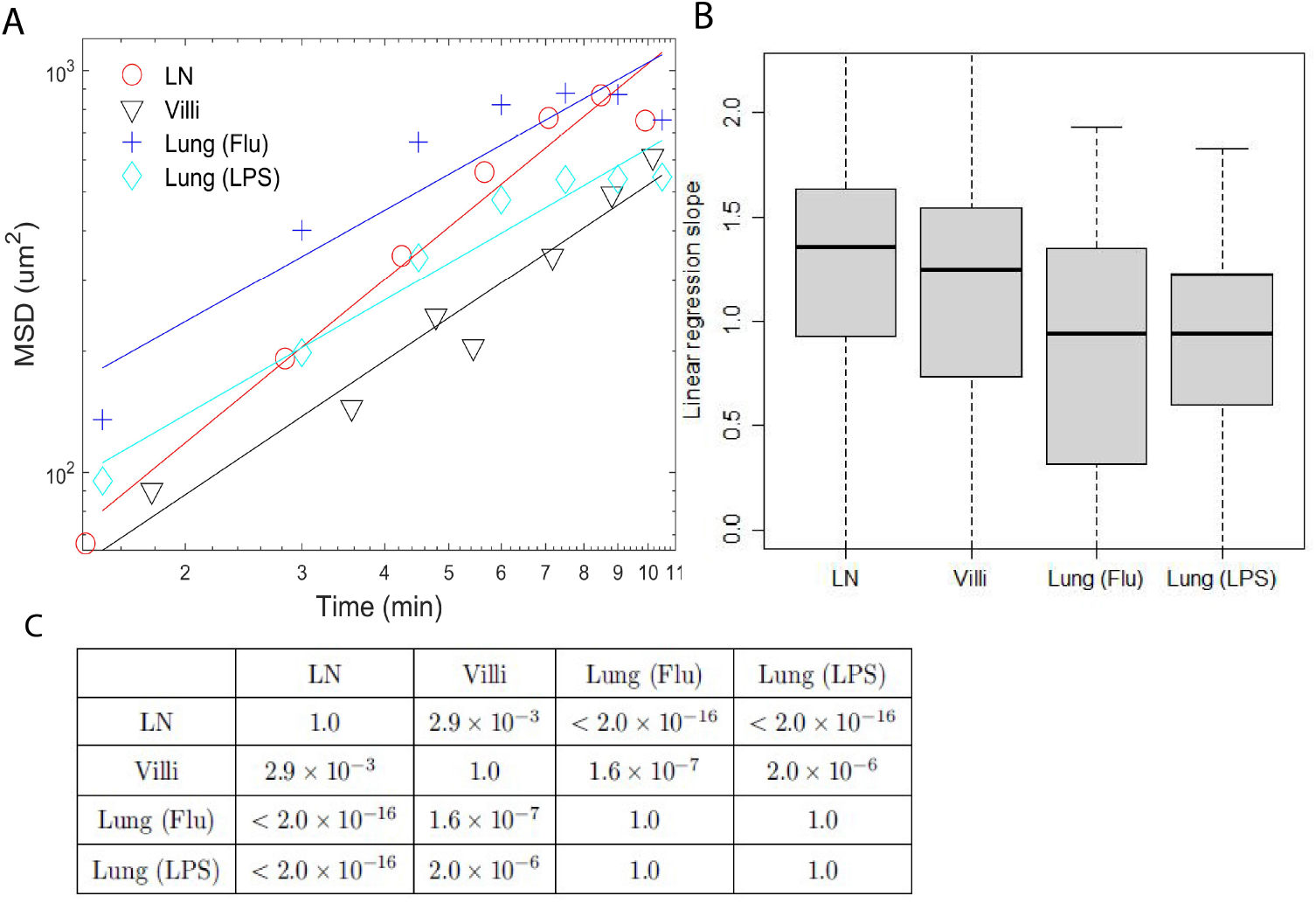
MSD of reduced data set. (A) Plots of mean square displacement (MSD) vs time and linear regression lines of individual representative cells near median from Figure 2B - supplement 2. (B) Box-and-whisker plots of linear regression cell slopes of log transformed mean squared displacement vs time. The median values are LN (1.35), villi (1.2), lung (Flu) (0.93), and lung (LPS) (0.94). (C) Table showing mean square displacement p-values as pairwise comparisons from Figure 2B - supplement 2 using Wilcoxon rank sum test.

**Figure 3 - supplement 2:**
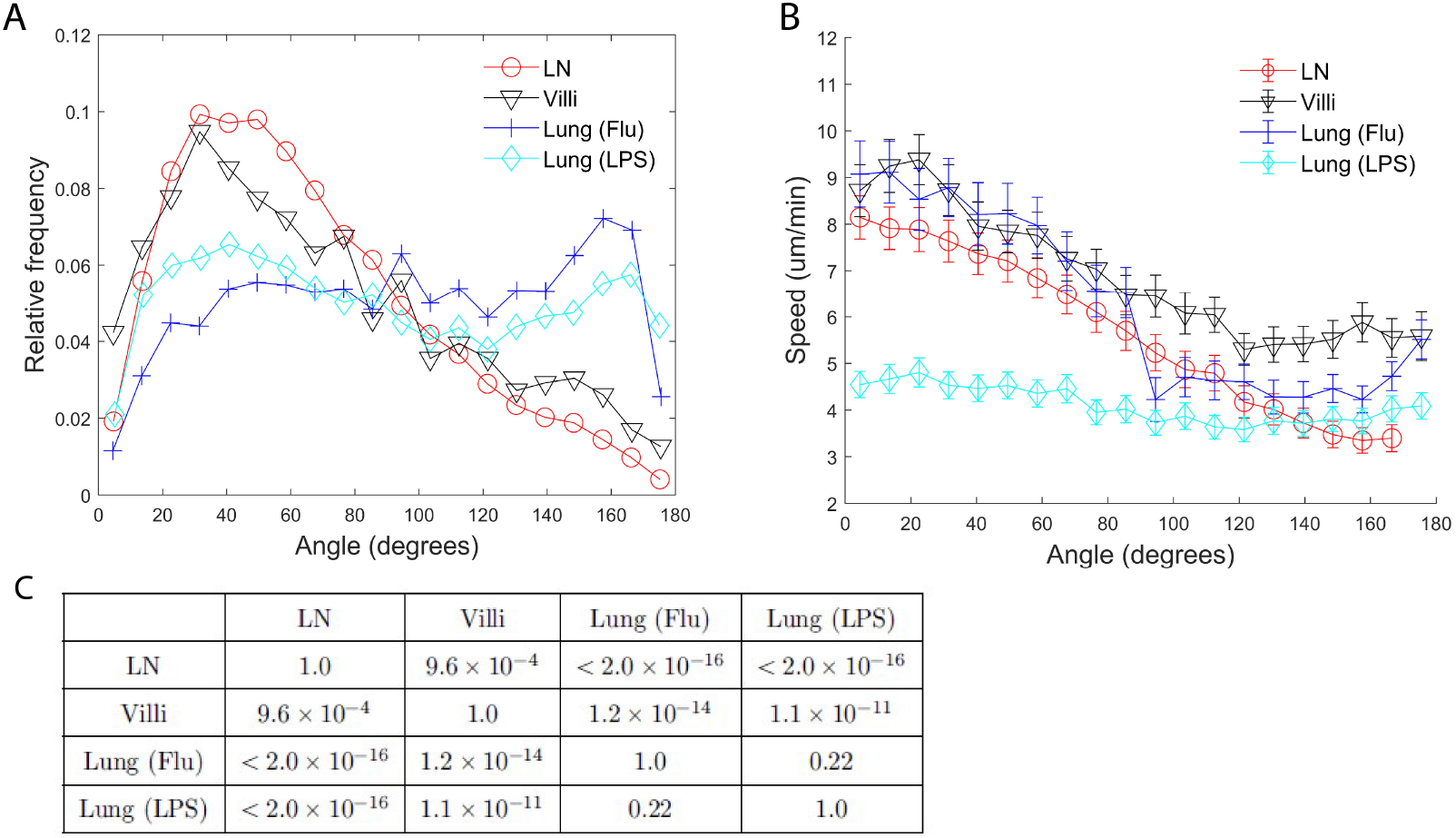
Turning angles and coupling of speed and turning angles of T cells from the reduced data set. (A) Relative frequency distribution of turning angles in each tissue. (B) Plot of speed (um/minute) versus angle (degrees). The speed tends to decrease as the turning angle increases in all tissues except for T cells in the LPS-inflamed lung. Error bars show plus and minus 1/8 of the standard deviation within each 9*^o^* angle bin. (C) Table of p-values of pairwise comparisons of proportion of turning angles less than 90*^o^* shown in Figure 3A - supplement 2 using Wilcoxon rank sum test.

**Figure 4 - supplement 2:**
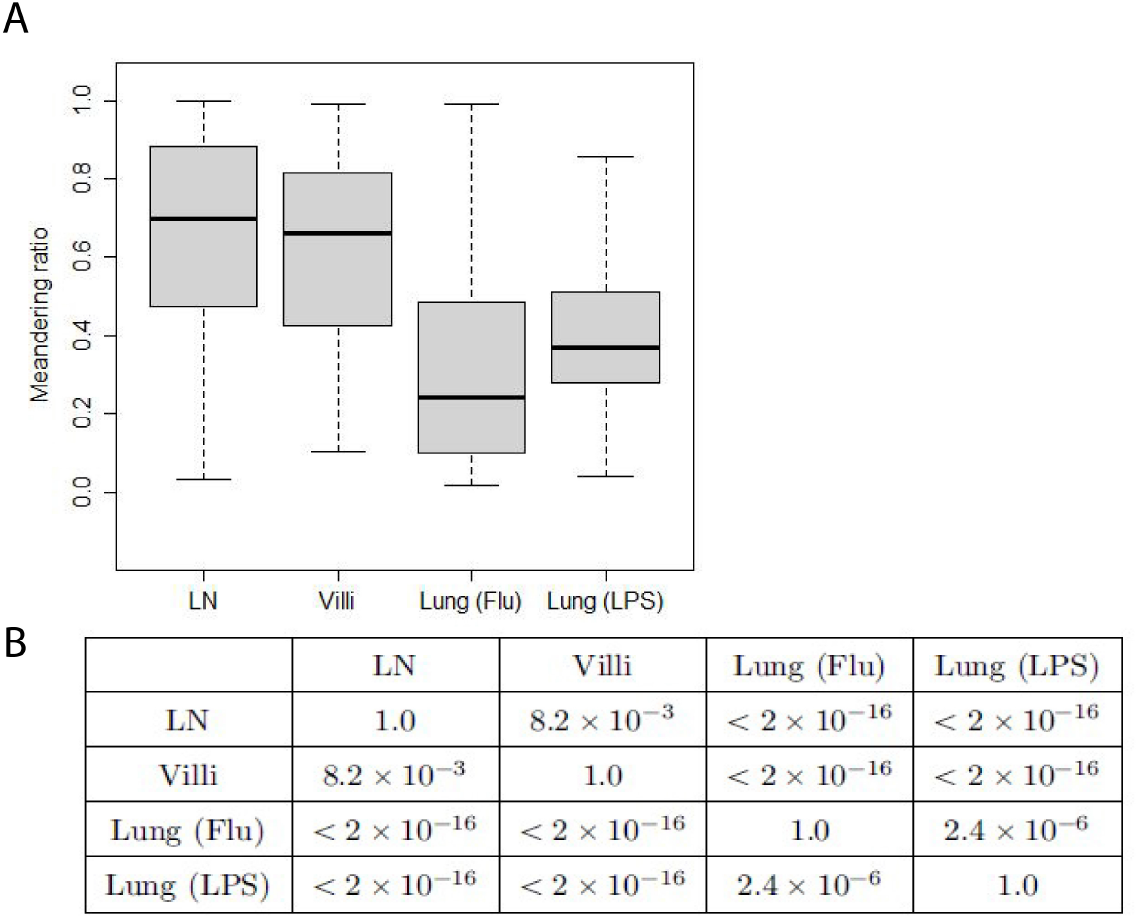
(A) Meandering ratio from the reduced data set. The median values are LN (0.70), villi (0.66), lung (Flu) (0.24), and lung (LPS) (0.37). (B) Table shows p-values of pairwise comparisons of meandering ratio as shown in Figure 4 - supplement 2 using Wilcoxon rank sum test.

**Figure 5 - supplement 2:**
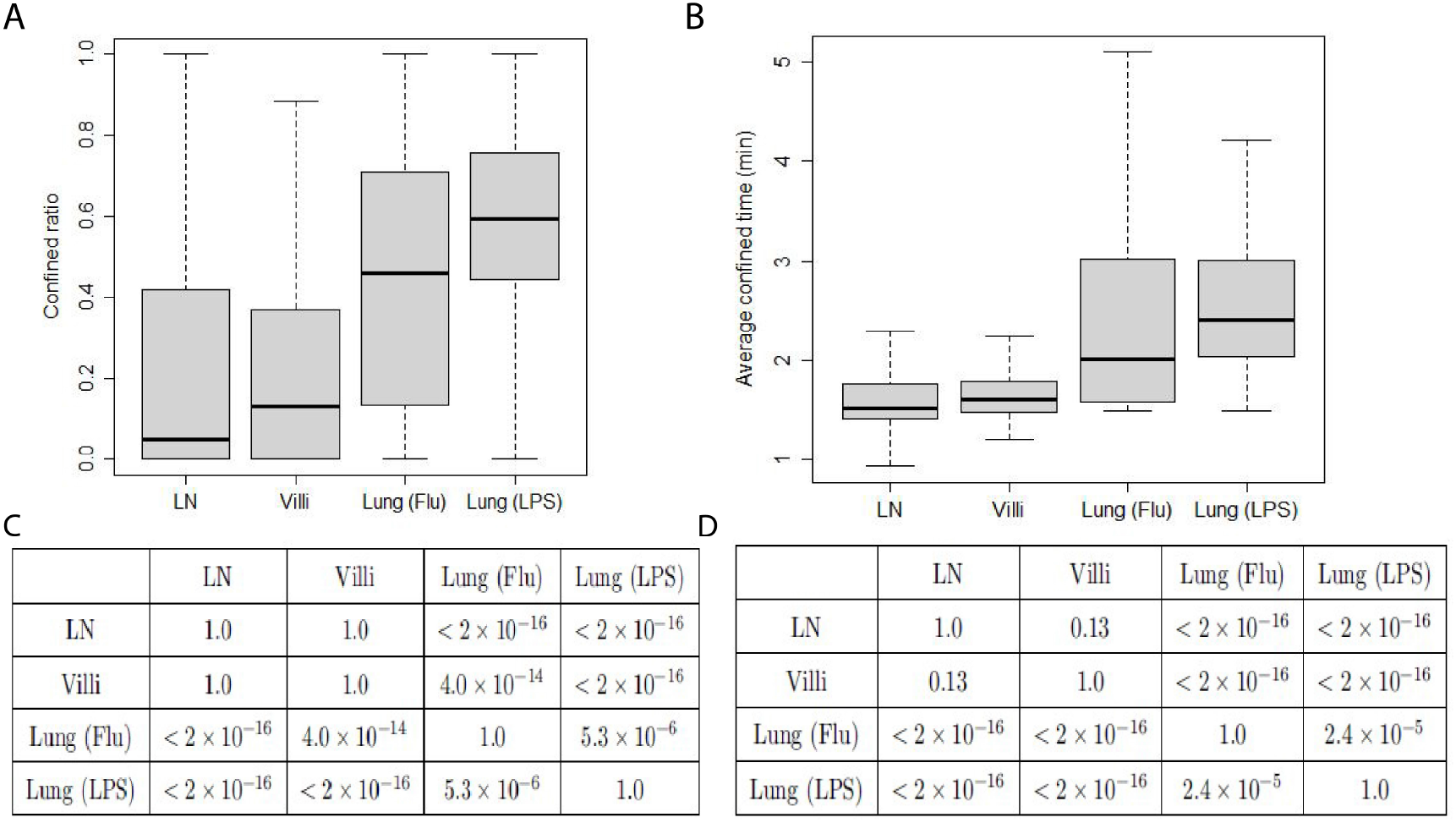
Confinement of T cells from the reduced data set. (A) Box-and-whisker plot of confined ratios. Median values: LN 0.05, Villi 0.13, Lung (Flu) 0.46, Lung (LPS) 0.60. (B) Box-and-whisker plot of confined time. Median values (minutes): LN 1.5, Villi 1.6, Lung (Flu) 2.0, Lung (LPS) 2.4. (C) Table showing p-values of pairwise comparisons of confined ratios from Figure 5A - supplement 2 using Wilcoxon rank sum test. (D) Table showing p-values of pairwise comparisons of confined time from Figure 5B - supplement 2 using Wilcoxon rank sum test.

**Figure 6 - supplement 2:**
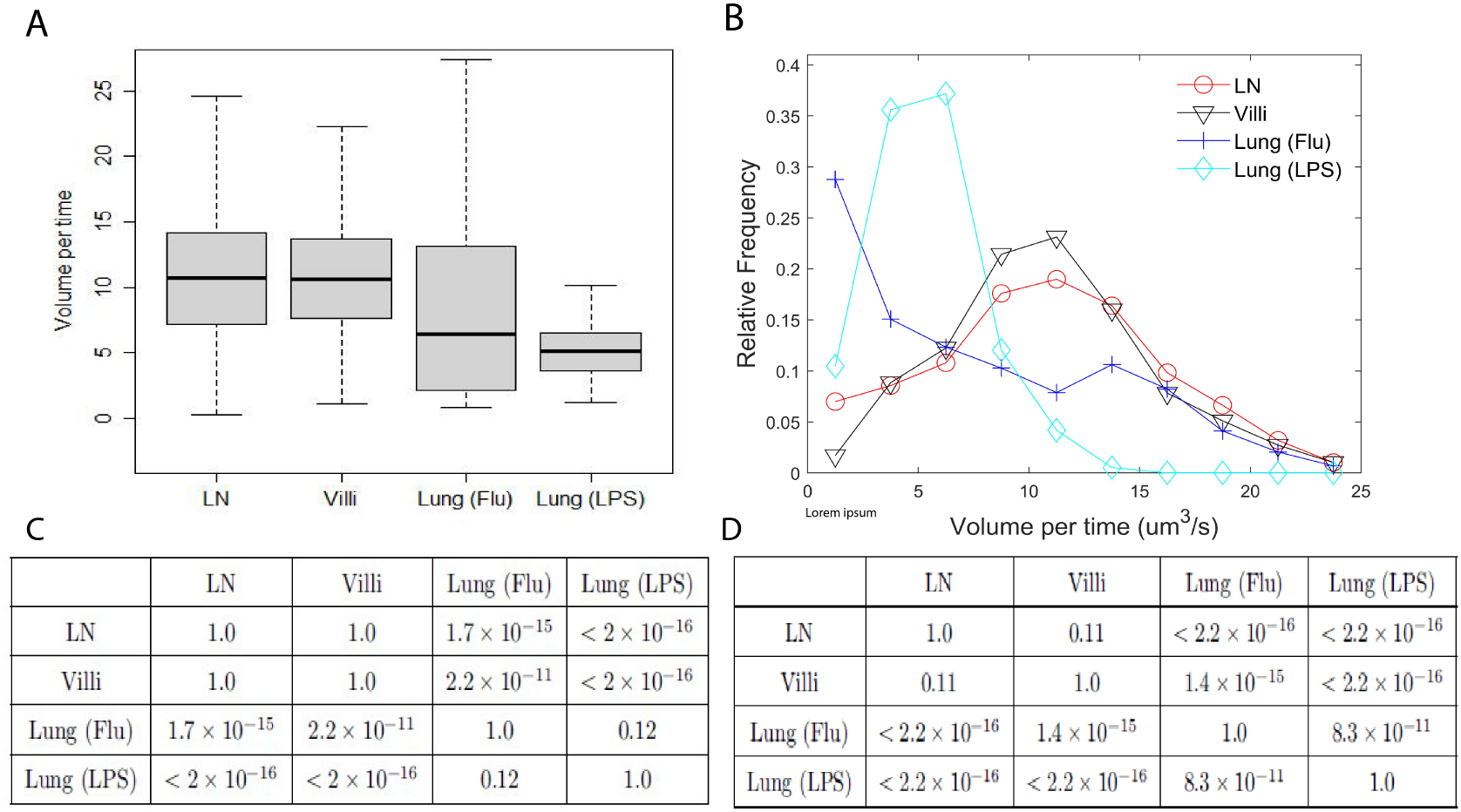
Volume patrolled by T cells in the reduced data set. (A) Box-and-whisker plot of median volume per time (cubic microns per second) patrolled by T cells in LN (10.7), villi (10.6), lung (flu) (6.4), and lung (LPS) (5.1). (B) Relative frequency distribution of volume per time (cubic microns per second) patrolled by T cells in each tissue. (C) Table of p-values of pairwise comparisons of volume per time from Figure 6A - supplement 2 using Wilcoxon rank sum test. (D) Table of p-values of pairwise comparisons of volume per time distributions from Figure 6B - supplement 2 using Kolmogorov-Smirnov test.

## Video legends

Video 1: GFP+ naïve T cells moving in lymph nodes. Naïve T cells were isolated from lymph nodes and spleen of Ubiquitin-GFP animals and adoptively transferred into naïve C57Bl/6 recipients, then imaged using 2 photon microscopy as described in Fricke et al. 2016 PLoS Computational Biology. GFP+ naïve T cells were imaged as described, tracked, and analyzed. The video contains a representative image from multiple fields of LNs imaged. The data is reproduced under the Creative Commons CC-BY 4.0 license.

Video 2: Migration of CD8 T Cells in Small Intestinal Villi at Day 8 after LCMV Infection. Naive P14-GFP CD8 T cells were transferred to B6 mice that were infected with LCMV one day later. At days 5 and 8 after infection, the jejunum was imaged via TPLSM. The representative time-lapse videos show P14-GFP CD8 T cells (cyan) at the indicated time points. Hoechst stain (blue) was injected prior to imaging. Reproduced from Thompson et al. Cell Reports 2019 Video S1 under CC BY-NC-ND 4.0 license. Only D8 T cells are shown and D5 movie removed from original file.

Video 3: Migration of CD8+ effector T cells in explanted lungs from LPS-inflamed animals. Left panel: Maximum projection of movie-sequence capturing adoptively transferred T cells (green) within an explanted lung. Trajectories (white lines) show the position of analyzed cells over time. Hours:minutes:seconds are shown in the left bottom corner. Right panel: 3D depiction of cell positions (green circles) and trajectories (blue lines) over time. To improve depth perception, the image volume is rotating during replay. Reproduced from Mrass et al. 2017 Nature Communications Supplementary Movie 1 under Creative Commons CC-BY 4.0 license.

Video 4: Migration of CD8+ effector T cells in lungs at d8 after HKx31 influenza infection. Naïve CD8 T cells from Ubiquitin-GFP animals were isolated and adoptively transferred into naïve C57Bl/6 mice, then infected with 1 × 10^3^ HKx31. At d8 post infection, lungs were removed and imaged with a heated and oxygenated chamber.

## Notes

https://github.com/davytorres/T-cell-analysis-tool

### Competing Interest Statement

The authors have declared no competing interest.

### Summary of Updates

The main manuscript has been revised so the Day 8 T cells from villi are used in the analysis instead of Day 5 T cells from the villi to be consistent with the days used in the other tissues. An ANOVA statistical analysis is also done to compare between tissue and within tissue variability.

